# Dephosphorylation of the pre-initiation complex during S-phase is critical for origin firing

**DOI:** 10.1101/2021.11.02.466916

**Authors:** Fiona Jenkinson, Kang Wei Tan, Barbara Schöpf, Miguel M Santos, Ivan Phanada, Philip Zegerman

## Abstract

Genome stability requires complete DNA duplication exactly once before cell division. In eukaryotes, cyclin-dependent kinase (CDK) plays a dual role in this regulation by inhibiting helicase loading factors before also activating origin firing. CDK activates initiation by phosphorylation of two substrates, Sld2 and Sld3, forming a transient and limiting intermediate – the pre-initiation complex (pre-IC). The importance and mechanism of dissociation of the pre-IC from origins is not understood. Here we show in the budding yeast *Saccharomyces cerevisiae* that CDK phosphorylation of Sld3 and Sld2 is specifically and rapidly turned over during interphase by the PP2A and PP4 phosphatases. Inhibiting dephosphorylation of Sld3/Sld2 causes dramatic defects in replication initiation genome-wide, retention of the pre-IC at origins and cell death. These studies not only provide a mechanism to guarantee that Sld3 and Sld2 are dephosphorylated before helicase loading factors but also uncover a novel positive role for phosphatases in eukaryotic origin firing.

## Introduction

Eukaryotes replicate their genomes from multiple origins that must initiate only once during the cell cycle. This is achieved by complete separation of the DNA loading of the replicative Mcm2-7 helicase (also known as licensing) and the activation of this helicase into different phases of the cell cycle ^1^. Cyclin-Dependent Kinase (CDK) plays a vital dual role in this regulation both as an inhibitor of licensing and together with Dbf4-dependent kinase (DDK) as an activator of the helicase in S-phase ^2^.

Although the exact details of how CDK and DDK mediate replication initiation is not fully understood, DDK directly phosphorylates Mcm2-7 double hexamers to generate a binding site for the initiation factors Sld3/Sld7, while CDK phosphorylates Sld3 and an additional initiation factor Sld2, which results in their phospho-dependent interaction with the BRCT repeats of Dpb11 ^2^. This complex formed by DDK and CDK on Mcm2-7 double hexamers is called the pre-initiation complex (pre-IC) and forms transiently at origins during the initiation reaction ^3, 4^. As Sld2, Sld3 and Dpb11 are not part of the replisome, they must be released by some mechanism during origin firing ^2^. It is not currently known whether the release of Sld2, Sld3 and Dpb11 is in itself an important step in the assembly of the replisome.

To ensure the complete replication of the entire genome, the number and timing of replication initiation events across chromosomes is also regulated during S-phase in eukaryotes ^5^. Although factors such as chromatin context determine the timing and likelihood of origin firing in S-phase ^5^, we and others have shown that key components of the pre-IC, including Sld2, Sld3 and Dpb11 as well as the Dbf4 subunit of DDK are stoichiometrically low in abundance and limit the number of simultaneous initiation events ^6–8^. This suggests that the active removal of the pre-IC from origins may be necessary to release and recycle these low abundance factors for further origin firing in a single S-phase (Figure 1A).

**Figure 1.**
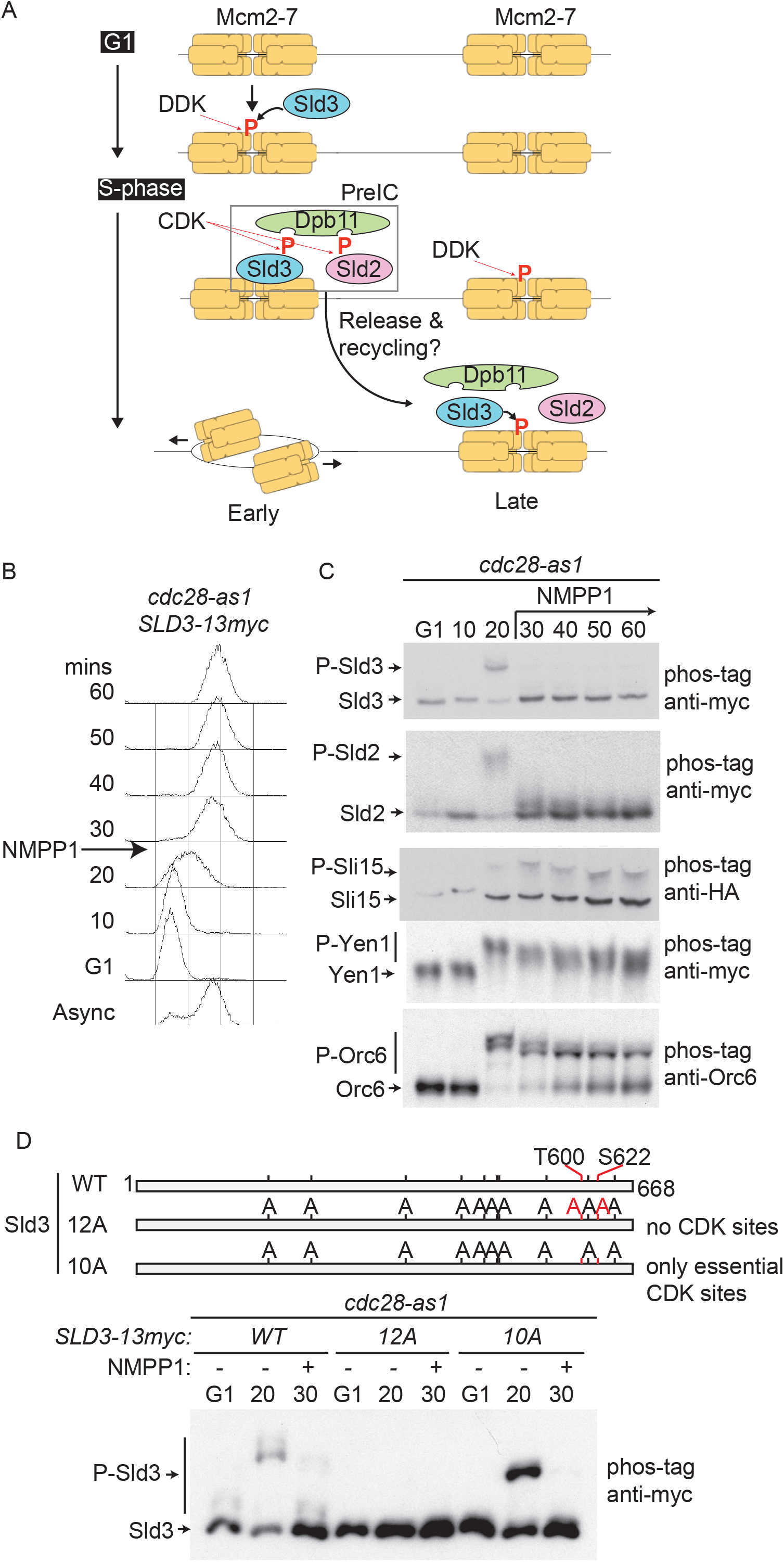
Sld2 and Sld3 are rapidly and selectively dephosphorylated in S-phase in *vivo*. A. Schematic diagram of pre-IC formation at an early origin (left) before a late origin (right). Many proteins are omitted for simplicity. B. Flow cytometry profile of the indicated strain arrested in G1 phase with alpha factor and released into S-phase for the indicated time. The *cdc28-as1* inhibitor 1-NM-PP1 was added at 25 minutes. 1-NM-PP1 is abbreviated in all figures to NMPP1. C. Phos-tag SDS page western blots of the indicated tagged strains using the cell synchronisation strategy from B. D. Top: scale diagram of Sld3, with the 10 non-essential CDK sites indicated in black and the 2 essential CDK sites T600 and S622 in red. WT stands for wild type. The 12A mutant has all CDK sites mutated to alanine, the 10A mutant still retains the essential CDK sites. Bottom: Block and release of the indicated strains as in B. 1-NM-PP1 was added at 25 minutes. These Sld3 alleles are expressed as a second copy.

Here we set out to analyse the mechanism of pre-IC regulation during S-phase in the budding yeast *Saccharomyces cerevisiae*. We show that the two critical CDK targets Sld2 and Sld3 are actively and specifically dephosphorylated during S-phase. The counteraction of Sld3/Sld2 phosphorylation is mediated by the PP2A and PP4 phosphatases and failure to dephosphorylate these pre-IC factors results in dramatic defects in replication initiation *in vivo*. Together we show that beyond the known role for kinases in origin firing, the dephosphorylation and dissolution of the pre-IC is a novel critical step in the eukaryotic replication reaction.

## Results

### Sld2 and Sld3 are rapidly and specifically dephosphorylated in S-phase

As the CDK-dependent pre-IC complex is a transient intermediate during origin firing, we hypothesised that a phosphatase might be required for the dissolution of this complex during S-phase (Figure 1A). To test this in budding yeast we inhibited CDK specifically in S-phase by synchronising cells containing the analogue sensitive (as) allele of the CDK catalytic subunit Cdc28, which is rapidly inhibited by the addition of 1-NM-PP1 ^9^ (Figure 1B). Analysis of CDK targets, Sld3, Sld2, Sli15, Yen1 and Orc6 revealed that these proteins are hypo-phosphorylated in G1 phase and phosphorylated as CDK levels rise on entry into S-phase (20 minutes, Figure 1C), as expected. Importantly, after addition of 1-NM-PP1 at 25 minutes, both Sld2 and Sld3 were rapidly dephosphorylated, but other CDK targets remained phosphorylated (Figure 1C). Unlike the rapid dephosphorylation of Sld2/Sld3, the slow accumulation of unphosphorylated Orc6 (Figure 1C) was reduced by cycloheximide treatment suggesting that this is due to new protein synthesis (Supplementary Figure 1A).

Sld3 has 12 CDK sites, two of which are essential for replication initiation, T600 and S622 ^10, 11^. To determine whether the rapid dephosphorylation of Sld3 after inhibition of CDK occurs at these essential sites we analysed the phosphorylation of Sld3 mutants that either have all CDK sites mutated to alanine (12A), or just the 10 non-essential sites (10A, Figure 1D). As expected, the wild-type protein was phosphorylated in S-phase and dephosphorylated upon addition of 1-NM-PP1, while the 12A mutant was not detectibly phosphorylated (Figure 1D). Significantly, the 10A mutant, which retains the essential CDK sites, was still phosphorylated and rapidly dephosphorylated upon addition of 1-NM-PP1 (Figure 1D), suggesting that the essential CDK sites in Sld3 that are required for replication initiation are rapidly dephosphorylated in S-phase in the absence of CDK.

Previous studies have shown that phosphorylation of Sld2 at multiple CDK sites allows phosphorylation of the CDK site (T84) required for binding to Dpb11 ^12^. Consistent with this co-dependence, any combination of mutations in the CDK sites of Sld2 abrogated all detectible phosphorylation (Supplementary Figure 1B-C), which prevented us from narrowing down which CDK sites in Sld2 are specifically dephosphorylated *in vivo*. Despite this, these data show that the CDK phosphorylation of both Sld3 and Sld2 are rapidly reversed in S-phase if CDK is inhibited.

### Sld2 and Sld3 are specifically dephosphorylated throughout S- and G2-phase independently of Rif1 and Cdc14

Yeast strains that lack the cyclin Clb5 have a temporal gap in CDK activity in S-phase ^13^ and in accordance with Figure 1 we wondered whether this gap would result in Sld2/Sld3 dephosphorylation. As expected, both Sld2 and Sld3 became phosphorylated in early S-phase in the *clb5Δ* mutant strain (20 minutes, Figure 2A), likely due to the activity of the alternative S-phase cyclin Clb6. Importantly, by 30 minutes both Sld2 and Sld3, but not Orc6, became dephosphorylated (Figure 2A). Interestingly, Sld2 phosphorylation increased again from 40 minutes, perhaps due to the accumulation of mitotic cyclin-CDKs, but Sld3 did not, suggesting different specificities of the CDK complexes (Figure 2A). This rapid and specific dephosphorylation of the key initiation factors Sld2/Sld3 in mid-S-phase may explain why only early origins initiate in *clb5Δ* strains ^14^.

**Figure 2.**
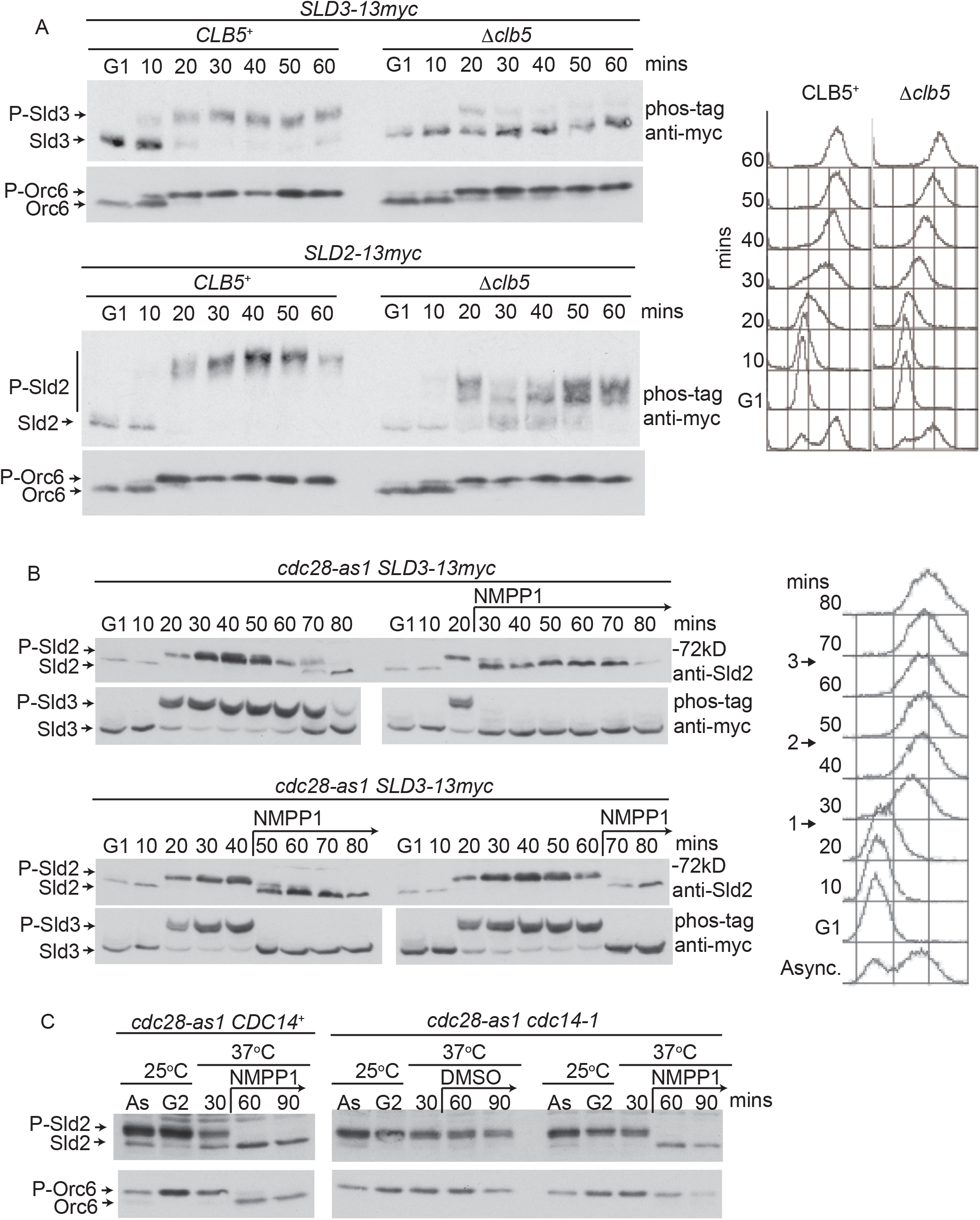
Sld2 and Sld3 are dephosphorylated throughout interphase independently of Cdc14. A. Left: Western blots from the indicated strains, blocked in G1 phase and released into S-phase. Right: Flow cytometry profile of the experiment on the left. B. Left: Western blots of Sld2 and Sld3 after the *cdc28-as1* inhibitor 1-NM-PP1 was either not added, or added at 25 (1), 45 (2) or 65 (3) minutes. Right: flow cytometry of the experiment on the right with 1, 2 or 3 indicating when 1-NM-PP1 was added. C. The indicated strains were arrested in nocodazole (G2) at 25°C and the temperature was shifted to 37°C for the indicated times, while still arrested in nocodazole. DMSO or 1-NM-PP1 was added at 30 minutes. Asynchronous cells are abbreviated as As.

To address the nature of this Sld2/Sld3 phosphatase, we explored when during the cell cycle this phosphatase activity was detectable. By synchronising the *cdc28-as1* strain in G1-phase and releasing into S-phase, we observed that addition of 1-NM-PP1 during early S-phase, late S-phase or G2 phase (Figure 2B) all resulted in rapid dephosphorylation of both Sld3 and Sld2, suggesting that the phosphatase activity is present during these periods of the cell cycle. Sld2 and Sld3 dephosphorylation was also not dependent on DNA replication as inhibition of helicase loading, using a temperature sensitive allele of Cdc6 (*cdc6-1*), prevented replication but not dephosphorylation (Supplementary Figure 2A).

It has been previously shown that the phosphatase required for mitotic exit in budding yeast, Cdc14, can dephosphorylate Sld2 and Orc6 *in vivo* ^15, 16^. Cdc14 is sequestered in the nucleolus until anaphase ^17^, suggesting that this is not the activity that is dephosphorylating Sld3/Sld2 from early S-phase onwards (Figure 2B). Despite this, some non-nucleolar activity has been detected for Cdc14 before anaphase ^18^. To test whether Cdc14 is responsible for Sld2/Sld3 dephosphorylation before mitotic exit, we utilised a temperature sensitive allele of Cdc14 (*cdc14-1*) to inhibit this phosphatase in nocodazole arrested cells. Importantly inhibition of *cdc14-1* prevented the dephosphorylation of Orc6, but not Sld2 or Sld3 (Figure 2C and Supplementary Figure 2B). This suggests that unlike Orc6, Sld2 and Sld3 are dephosphorylated independently of Cdc14 before mitosis.

The PP1 phosphatase, through binding to Rif1, has been shown to dephosphorylate DDK substrates ^5^. To address whether PP1/Rif1 might also be required for Sld2/Sld3 dephosphorylation we performed an experiment as in Figure 1C, but in a strain lacking Rif1. Significantly, while DDK-dependent Mcm4 phosphorylation was insensitive to inhibition of CDK, Sld2 and Sld3 were rapidly dephosphorylated in a *rif1Δ* strain (Supplementary Figure 2C). This again demonstrates the specificity of the targeted dephosphorylation of Sld2/Sld3 over other phosphorylated replication factors and shows that neither PP1/Rif1 nor Cdc14 are involved in Sld2/Sld3 dephosphorylation in S-phase.

### Sld2 and Sld3 are dephosphorylated in S-phase by PP2A and PP4

To narrow down the phosphatases responsible for Sld2/Sld3 dephosphorylation we used a chemical genetics approach (Supplementary Figure 3A). Two broad specificity phosphatase inhibitors, cantharidin and 9,10-phenanthrenequinone (PQ), appeared to abrogate Sld3 dephosphorylation *in vivo* (Supplementary Figure 3B). Sld3 is also phosphorylated by the checkpoint kinase Rad53 ^19, 20^, and while cantharidin did not cause Rad53 activation, PQ was a potent activator of Rad53 (Supplementary Figure 3C) likely due to oxidative DNA damage ^21^. Therefore, the hyperphosphorylation of Sld3 observed in PQ, but not cantharidin, is probably an artefact of Rad53 activation.

To confirm that cantharidin inhibits the dephosphorylation of both Sld3 and Sld2 we incubated the *cdc28-as1* strain with cantharidin, with and without the addition of 1-NM-PP1 (Supplementary Figure 3D). Addition of cantharidin for 10 minutes led to hyperphosphorylation of Sld3 even in the absence of 1-NM-PP1, while longer incubations resulted in the complete abrogation of the 1-NM-PP1 induced dephosphorylation of both Sld3 and Sld2 (Supplementary Figure 3D). Together these data show that cantharidin can fully counteract the rapid dephosphorylation of both Sld3 and Sld2. The rapid hyper-phosphorylation of Sld2/3 after inhibition of the phosphatase (cantharidin) and the rapid de-phosphorylation of Sld2/3 after inhibition of CDK (1-NM-PP1) demonstrate that Sld3 and Sld2 are dynamically and constantly phosphorylated and dephosphorylated in S/G2 phase (Supplementary Figure 3D/E).

Although cantharidin has a broad phosphatase inhibitory profile, it has the lowest IC_50_ for the PP2A family phosphatases (Supplementary Figure 4A and ^22^). In yeast, there are five PP2A family members, two of which, *PPH21* and *PPH22*, are close paralogues (Supplementary Figure 4A). While mutation of *SIT4* and *PPG1* had little effect on Sld3 dephosphorylation (data not shown), combined mutation of both *PPH21* and *PPH22* resulted in hyperphosphorylation and reduction of dephosphorylation of Sld3 and to a lesser extent of Sld2 (Figure 3A-B). Mutants of *pph21/pph22* have defects in budding and cytokinesis due to the activation of yeast Wee1 (Swe1) ^23^ and therefore strains with PP2A mutants are also *swe1Δ* to overcome these defects ^23, 24^. The loss of *SWE1* does not affect the dephosphorylation phenotypes we observe (e.g Figure 3D).

**Figure 3.**
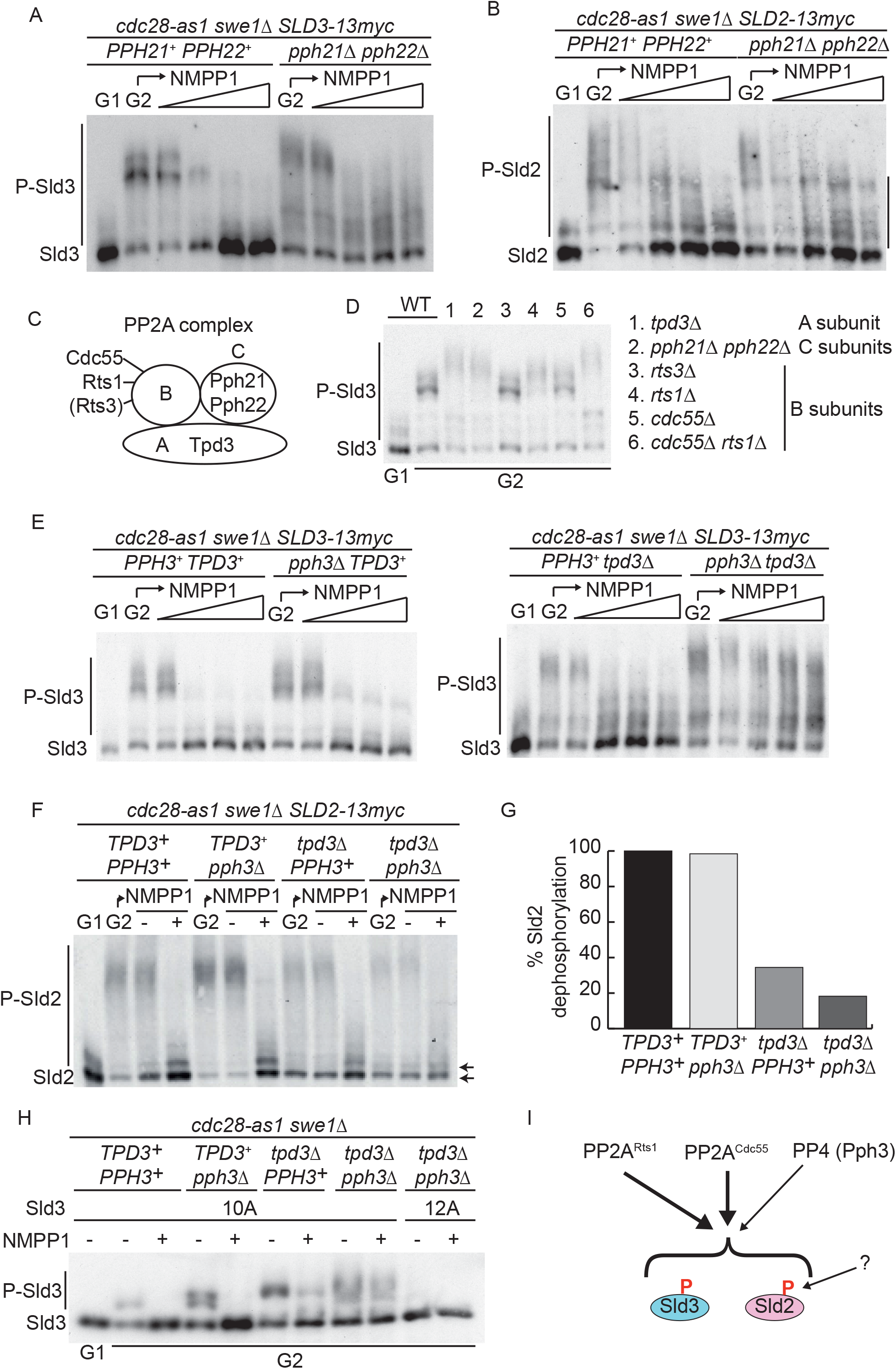
Identification of PP2A and PP4 (Pph3) as the Sld2 and Sld3 phosphatases *in vivo*. A. Western blot of Sld3-13myc from the indicated strains, arrested in nocodazole (G2) followed by the addition of an increasing concentration of 1-NM-PP1 up to 10μM for 10 minutes. A G1 arrested sample is loaded to indicate the hypo-phosphorylated protein. PP2A mutants have defects in budding and cytokinesis due to the activation of Swe1 ^23^ and therefore strains with PP2A mutants are also *swe1Δ* to overcome these defects, unless indicated. B. As A, but for Sld2-13myc. C. Schematic diagram of the budding yeast PP2A complexes. There are 2 paralogous catalytic (C) subunits (Pph21 and Pph22) and 2 alternative specificity (B) subunits (Cdc55 and Rts1), but only one scaffold (A) subunit (Tpd3). Rts3 is in brackets as it is only a putative component of PP2A. D. Western blot of Sld3-13myc from a wild type strain (WT) or the 6 indicated PP2A mutant strains arrested in nocodazole (G2). A G1 arrested sample is loaded to indicate the hypo-phosphorylated protein. These strains are *SWE1^+^*. E. As A. F. As B except nocodazole arrested cells (G2) were held for 10 minutes in the presence (+) or absence (−) of 5μM 1-NM-PP1. G. Quantification of the amount of the hypo-phosphorylated forms of Sld2 (arrows in F) after 1-NM-PP1 addition, relative to the *TPD3^+^ PPH3^+^* strain. H. As F, utilising the CDK site mutants of Sld3 in Figure 1D. I. Schematic diagram of the 3 phosphatase complexes that regulate Sld3 and Sld2. Residual dephosphorylation of Sld2 even in the *tpd3Δ pph3Δ* strain (see G) suggests there may be other phosphatase regulators of Sld2 (indicated by ?).

PP2A is a trimeric complex including a catalytic (C, Pph21/Pph22 – Figure 3C), scaffold (A, Tpd3) and a specificity (B) subunit (Cdc55 or Rts1)^25^. As with the *pph21Δ pph22Δ* strains, *tpd3Δ* resulted in hyper-phosphorylation of Sld3 (Figure 3D) and significantly reduced the dephosphorylation of Sld3 and to a lesser extent of Sld2 (Supplementary Figure 4B/C). Loss of the putative B subunit Rts3 had no effect on Sld3 phosphorylation (Figure 3D), whereas loss of either Rts1 or Cdc55 had a partial effect on Sld3 phosphorylation (Figure 3D) and loss of both mimicked the *tpd3Δ* and the *pph21Δpph22Δ* strains (Figure 3D and Supplementary Figure 4D) suggesting that PP2A complexes containing either Rts1 or Cdc55 both contribute to Sld3 dephosphorylation *in vivo*. As with the *pph21Δ pph22Δ* and *tpd3Δ* strains the effect of the *cdc55Δ rts1Δ* double mutant on Sld2 was less pronounced than for Sld3 (Supplementary Figure 4E).

Although PP2A mutants have a dramatic effect on Sld3 dephosphorylation, some dephosphorylation was still observed in the presence of 1-NM-PP1 (e.g Figure 3A), which was not the case after cantharidin treatment (Supplementary Figure 3D). It has been shown that in Pph21/Pph22 deficient cells another cantharidin-sensitive PP2A family phosphatase Pph3, which is the catalytic subunit of the PP4 complex, can contribute residual phosphatase activity ^26, 27^. To test whether Pph3 contributes to Sld2 and Sld3 dephosphorylation we combined null mutations of the PP2A scaffold subunit *tpd3Δ* with *pph3Δ*. Importantly while loss of Pph3 alone had a small effect on Sld3 or Sld2 dephosphorylation (Figure 3E and 3F) the *tpd3Δ pph3Δ* double mutant completely lacked any Sld3 dephosphorylation in the presence of 1-NM-PP1 (Figure 3E, right panel), similar to cantharidin (Supplementary Figure 3D). This phosphorylation of Sld3 was not due to spurious Rad53 activation (Supplementary Figure 4F). For Sld2, we still observed some dephosphorylation in the *tpd3Δ pph3Δ* double mutant (Figure 3F), but quantification of the hypo-phosphorylated forms of Sld2 revealed that the dephosphorylation of Sld2 was dramatically reduced in the *tpd3Δ pph3Δ* strain (Figure 3G).

PP2A and Pph3 dephosphorylate the essential CDK sites of Sld3 because the 10A mutant, which only retains the 2 essential CDK sites (Figure 1D), was not dephosphorylated in the *pph3Δ tpd3Δ* double mutant strain (Figure 3H). Together, these data show that both PP2A^RTS1^ and PP2A^CDC55^, but also PP4 (Pph3) are responsible for the dephosphorylation of the CDK sites in Sld3 and also in large part for the dephosphorylation of Sld2 (Figure 3I).

### Sld3 interaction with Rts1 contributes to dephosphorylation

Having established that PP2A and Pph3 control Sld3/Sld2 dephosphorylation in S-phase we wanted to determine the physiological role of this regulation. Unfortunately, due to the large number of functions of these phosphatases, combined mutations of PP2A and Pph3 are extremely sick ^26, 27^. To examine specifically the functional importance of Sld3/Sld2 dephosphorylation we set out to identify phosphatase-interaction mutants of Sld2/Sld3 that would be defective only in the regulation of these CDK targets. A short linear motif (SLiM) has been identified that is required for the recruitment of the mammalian orthologue of Rts1 (B56) to substrates ^28, 29^. Using the B56 binding-site prediction software ^29^, we identified a high probability Rts1 binding site (LxxIxE) in Sld3, starting at amino acid 616 (Figure 4A and Supplementary Figure 5A). Interestingly, this LxxIxE motif is immediately followed by one of the essential CDK sites in Sld3, S622 (Figure 4A).

**Figure 4.**
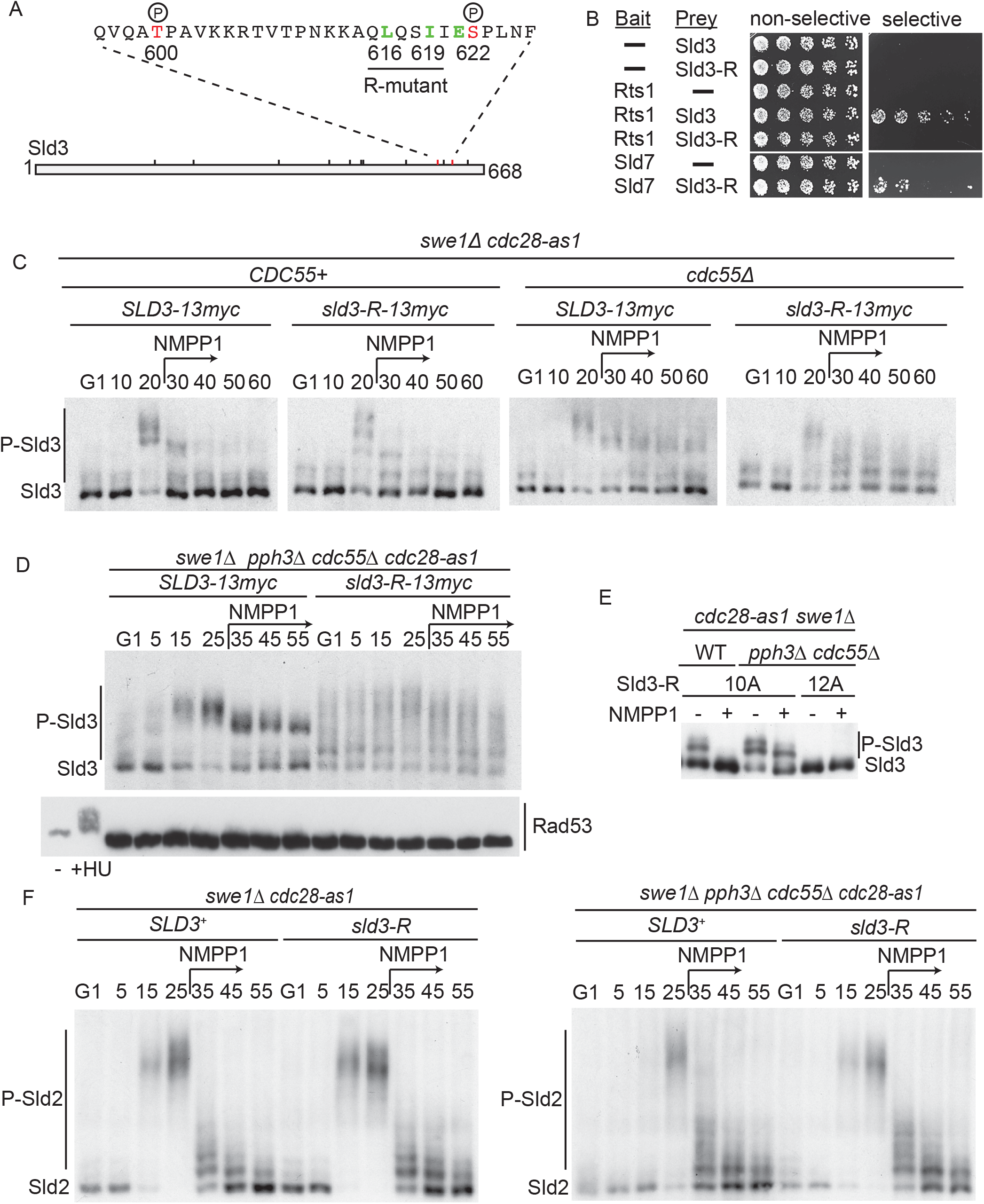
Sld3 is targeted by PP2A^RTS1^ through interaction with Rts1. A. Schematic diagram of Sld3 with the sequence of the region of the essential CDK sites above. The putative Rts1 binding consensus sequence LxxIxE is indicated in green. B. Yeast two-hybrid analysis of the indicated proteins. – indicates empty bait/prey plasmid. Sld3-R refers to the Sld3 protein with 2 residues of the Rts1 binding consensus (L616 and I619) mutated to alanine. C. Phos-tag anti-myc western blot of the indicated strains arrested in G1 and released into S-phase. 1-NM-PP1 was added at 25 minutes D. As C, except 1-NM-PP1 was added at 30 minutes. E. As Figure 3H. F. As D, but for Sld2-13myc.

Sld3 indeed interacts with Rts1 by yeast-two-hybrid analysis (Figure 4B), and importantly mutation of the Rts1-SLiM in Sld3 (L616A and I619A), hereafter called the *sld3-R* allele, abrogated Rts1 binding (Figure 4B). The L616A and I619A mutations did not affect the binding of Sld3 to other replication factors, including Sld7 and Dpb11 (Figure 4B and Supplementary Figure 5B). As a result, the *sld3-R* allele is viable and is not synthetic lethal with other replication or checkpoint mutants (Supplementary Figure 5C).

Having identified an Rts1 binding motif in Sld3, we set out to establish whether this interaction is important for the dephosphorylation of Sld3 *in vivo*. Similar to the null mutations in Rts1 (Figure 3D), the *sld3-R* mutant showed a minor defect in Sld3 dephosphorylation (Figure 4C) but combination of the *sld3-R* mutant with *cdc55Δ* or *pph3Δ* increased the defect in dephosphorylation (Figure 4C and Supplementary Figure 5D), while a triple mutant *sld3-R cdc55Δ pph3Δ* completely abrogated the dephosphorylation of Sld3 in S-phase (Figure 4D). This triple mutant that lacks the interaction with PP2A^RTS1^ and lacks PP2A^CDC55^ and PP4 (Pph3) also resulted in abundant phosphorylation of Sld3 in G1 phase (Figure 4D), perhaps due to CDK phosphorylation remaining from previous cycles (see Discussion). Importantly, loss of Sld3 dephosphorylation in the *sld3-R cdc55Δ pph3Δ* strain was not due to inappropriate Rad53 activation (Figure 4D), occurred at the essential CDK sites (Figure 4E) and did not affect Sld3 protein stability (Supplementary Figure 5F). Together these data show that we have identified a separation of function mutant of Sld3 that can no longer be targeted by PP2A^RTS1^ and in combination with mutation of PP2A^CDC55^ and PP4 results in abrogation of the dephosphorylation of Sld3 in S-phase.

Since Sld2 is also dephosphorylated in a PP2A-dependent manner (Figure 3F/G) we wondered whether the Rts1-binding mutant of Sld3 might affect Sld2 phosphorylation *in trans*. Although fully defective in Sld3 dephosphorylation (Figure 4D), the *sld3-R cdc55Δ pph3Δ* strain was only partially defective in Sld2 dephosphorylation (Figure 4F and Supplementary Figure 5E).

### Failure to dephosphorylate Sld3 and Sld2 causes a defect in origin firing

To specifically test for S-phase defects resulting from the failure to dephosphorylate Sld2/Sld3 we utilised conditional mutants of *CDC55* and *PPH3*. Auxin-inducible degrons (AID) of Cdc55 and Pph3 showed specific degradation in the presence of auxin (Supplementary Figure 6A-B), as expected ^30, 31^. Timely inhibition of *cdc55-AID* and *pph3-AID* in G1 phase combined with the *sld3-R* mutant resulted in abrogation of the dephosphorylation of Sld3 during the subsequent S-phase (Supplementary Figure 6C), but importantly largely prevented the hyper-phosphorylation of Sld3 in G1 phase observed in the null mutants (compare Supplementary Figure 6C with Figure 4D). This suggested that the conditional degron alleles of Cdc55 and Pph3 combined with the *sld3-R* mutant is a suitable inducible strain to analyse specifically the S-phase functions of the dephosphorylation of Sld3/Sld2.

Flow cytometry analysis revealed that failure to dephosphorylate Sld3 (*cdc55-AID pph3-AID sld3-R*) resulted in a dramatically slower S-phase (purple versus grey, Figure 5A and 5B), which was not due to defects in the G1-S transition (as determined by budding index, Figure 5C), nor due to aberrant Rad53 activation (Figure 5D). To analyse this defect in S-phase progression in more detail we determined the dynamics of DNA replication genome-wide by high-throughput sequencing and copy-number analysis. Plotting the median replication time (T_rep_) produces a profile whereby peaks delineate origins and troughs represent termination zones (Figure 5E, Chromosome XIV is shown as an example). While the control strain revealed peaks of early and later origin firing (e.g ARS1415 and ARS1411 respectively, black line Figure 5E), the *cdc55-AID pph3-AID sld3-R* mutant that cannot dephosphorylate Sld3 showed a delay in early origin firing and a dramatic defect in late origin firing (purple line, Figure 5E). Significantly we did not detect any difference in replication elongation rates (as determined by the slope of T_rep_ between origins), suggesting that the slow S-phase in the *cdc55-AID pph3-AID sld3-R* strain is due to initiation not replisome progression defects (data not shown).

**Figure 5.**
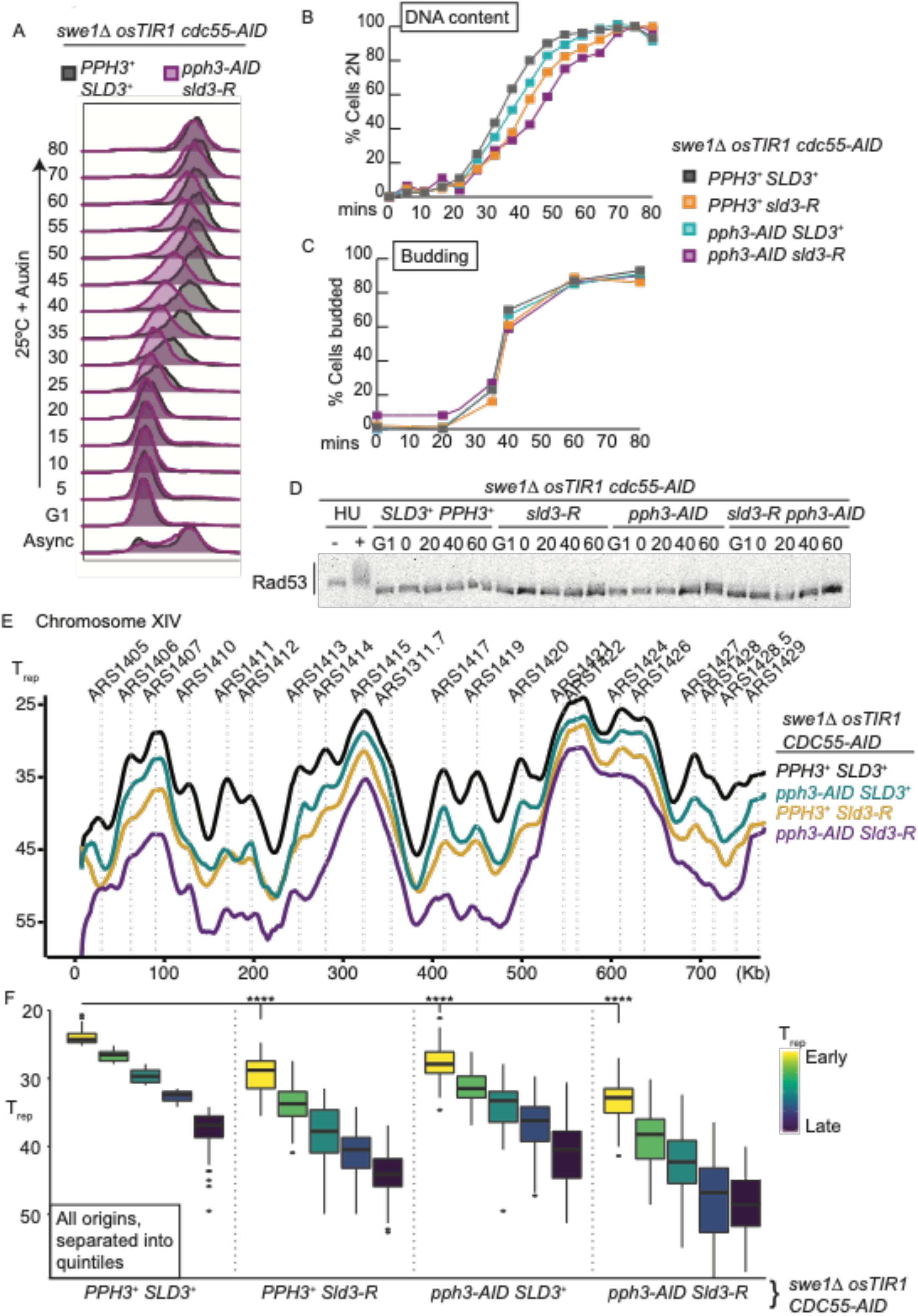
Abrogation of Sld3/2 dephosphorylation in S-phase causes defects in origin firing. A. Flow cytometry of the indicated strains arrested in G1 phase at 25°C, then held in G1 phase and shifted to 37°C in the presence of auxin for 30 minutes, before release into S-phase at 25°C in the presence of auxin. AID is the auxin inducible degron and osTir1 is required for degradation of this tag in the presence of auxin. B. DNA content (mean value of the histograms) from the flow cytometry in A. C. Budding index from A. D. Rad53 western blot from A. E. Replication profile of the indicated strains. T_rep_ is the median replication time. Only chromosome XIV is shown as an example, known origins are annotated above. F. Box and whisker plot of all origins from E, separated into quintiles according to their normal T_rep_. **** represents P value <10^−11^ from a t-test.

Analysis of all origins, binned into quintiles according to their normal T_rep_, revealed that every origin group was delayed in the *cdc55-AID pph3-AID sld3-R* strain, even the earliest firing origins (yellow boxes, Figure 5F), suggesting that dephosphorylation of Sld3 is required for all origin firing. Phosphatase mutant combinations with only a partial defect in Sld3 dephosphorylation also showed a partial defect in S-phase progression by flow cytometry (orange/blue lines, Figure 5B) and by copy number analysis (Figure 5E/F). These combinations reveal that the replication phenotype is not simply due to any one of the *cdc55-AID pph3-AID sld3-R* alleles having a defect in origin firing, but instead is an additive effect, similar to Sld3 dephosphorylation (e.g Figure 4C/D). Therefore, using the conditional Cdc55/Pph3 mutants in synchronised cells, combined with the separation of function allele of Sld3 that cannot bind to PP2A^RTS1^, Figure 5 demonstrates that dephosphorylation of Sld3/Sld2, is important for origin firing genome-wide.

### Failure to dephosphorylate Sld3/Sld2 delays the release of the pre-IC from origins

Since dephosphorylation of the pre-IC proteins Sld3/Sld2 is important for origin firing (Figure 5), we wondered whether this dephosphorylation is important for pre-IC release from origins (Figure 1A). Detection of the Mcm2-7 complex at origins through Mcm4 ChIP-seq (Figure 6A) showed that Mcm2-7 is loaded at all origins in G1 phase in both the control strain (*cdc55-AID*) and the strain that is defective in the dephosphorylation of Sld3/Sld2 (*cdc55-AID pph3-AID sld3-R)*. Importantly this demonstrates that the reduction in origin firing observed in the *cdc55-AID pph3-AID sld3-R* strain (Figure 5) is not due to a licensing defect. By 40 minutes after release from G1 phase the Mcm2-7 complex becomes delocalised from the earliest origins in the control strain, as initiation occurs (Figure 6A, note that the heatmaps are ordered from early to late firing origins). In the *cdc55-AID pph3-AID sld3-R* strain however, the movement of the Mcm2-7 complex away from origins was greatly delayed (Figure 6A), consistent with a reduction in initiation at all origins when Sld3/Sld2 dephosphorylation is defective (Figure 5).

**Figure 6.**
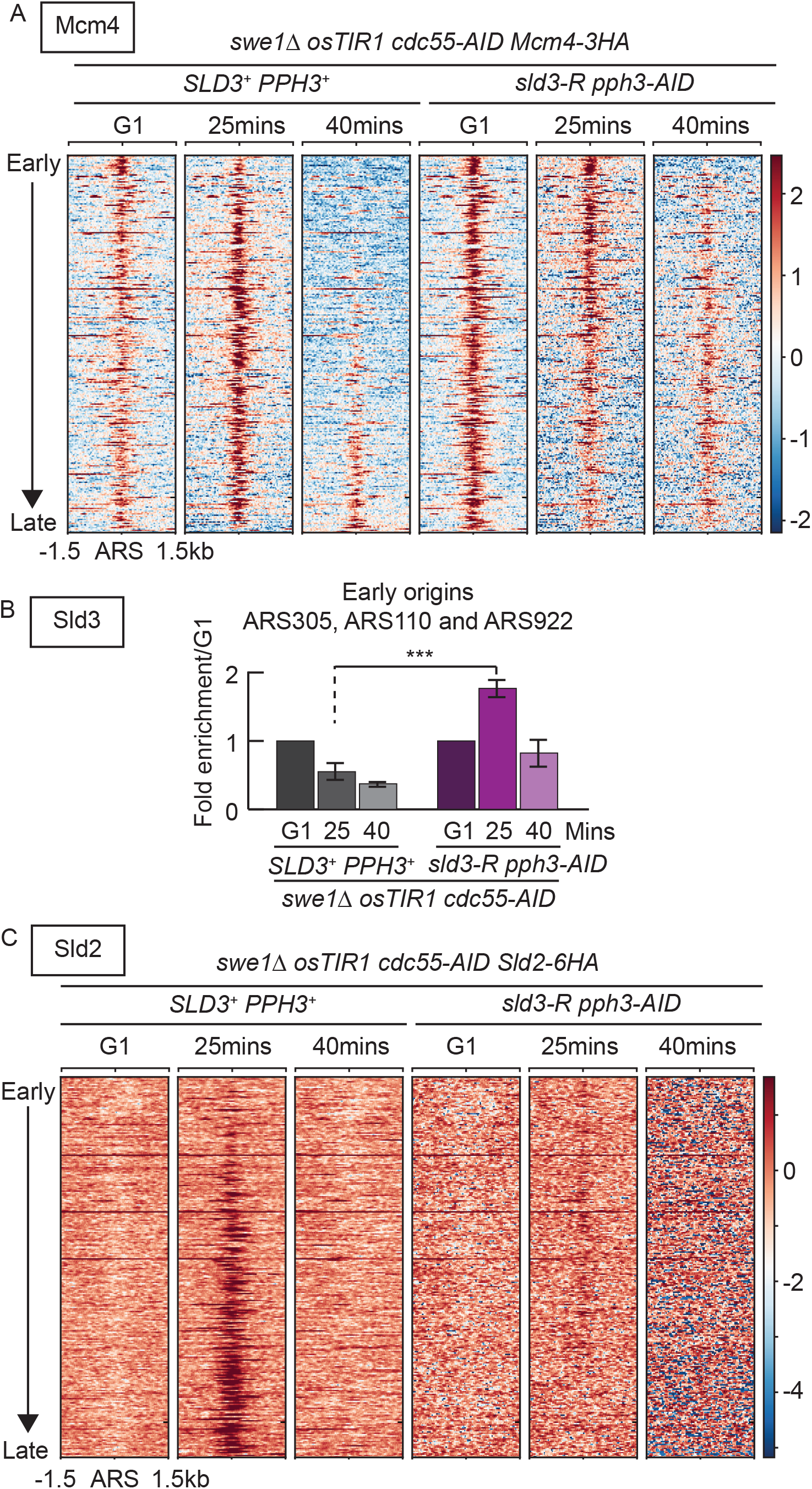
Abrogation of Sld3/2 dephosphorylation in S-phase causes the retention of the pre-IC at origins. A. Heatmap of Mcm4-3HA ChIP-seq form the indicated strains arrested in G1 phase and released into S-phase for 25 or 40 minutes. The heatmap is centred on known ARS elements, with 1.5kb of flanking sequence and arranged from top to bottom by increasing T_rep_. B. qPCR analysis of Sld3-13myc ChIP as in A, for the early origins ARS305, ARS110, ARS922. The G1 enrichment signal was set to 1, error bars are SD, n=4, *** is a P value of <0.0005 from a t-test C. As in A, but for Sld2-6HA

To analyse pre-IC dynamics we first performed Sld3 ChIP-seq but we could not achieve sufficient enrichment for Sld3 by this method (data not shown). Instead, we analysed Sld3-ChIP by qPCR, as previously described ^3^, at three early firing origins (Figure 6B). In the control strain, Sld3 binds to early origins in G1 phase and is released from origins during S-phase (Figure 6B). In the *cdc55-AID pph3-AID sld3-R* strain however, Sld3 accumulated at these origins during S-phase, relative to G1 (Figure 6B), consistent with a delayed release of this protein from origins in the absence of dephosphorylation. For Sld2, we observed a transient interaction with origins during S-phase as expected ^3^, which at 25 minutes was preferentially at later firing origins in the control strain (Figure 6C). In the *cdc55-AID pph3-AID sld3-R* strain Sld2 was still detected at early firing origins at 25 minutes (Figure 6C), reflecting the delay in initiation and delay in the release of Sld3 from early origins (Figure 6B). Sld2 is released from origins to a greater extent than Sld3 in the *cdc55-AID pph3-AID sld3-R* strain (Figure 6B versus 6C), likely because Sld2 is still dephosphorylated to a significant degree in this background (Figure 4F). Together these ChIP data are consistent with a role for dephosphorylation of Sld2/Sld3 in the release of the pre-IC from origins during initiation.

### Dephosphorylation of Sld3 and Sld2 is essential

Significantly, combination of *cdc55Δ*, *pph3Δ* and *sld3-R*, which abrogates Sld3/Sld2 dephosphorylation (Figure 4D/F) and causes genome-wide defects in replication initiation (Figure 5) was largely inviable, except for a small number of microcolonies (Figure 7A). This synthetic lethality was also observed with the *cdc55-AID pph3-AID sld3-R* strain in the presence of auxin (Supplementary Figure 6D).

**Figure 7.**
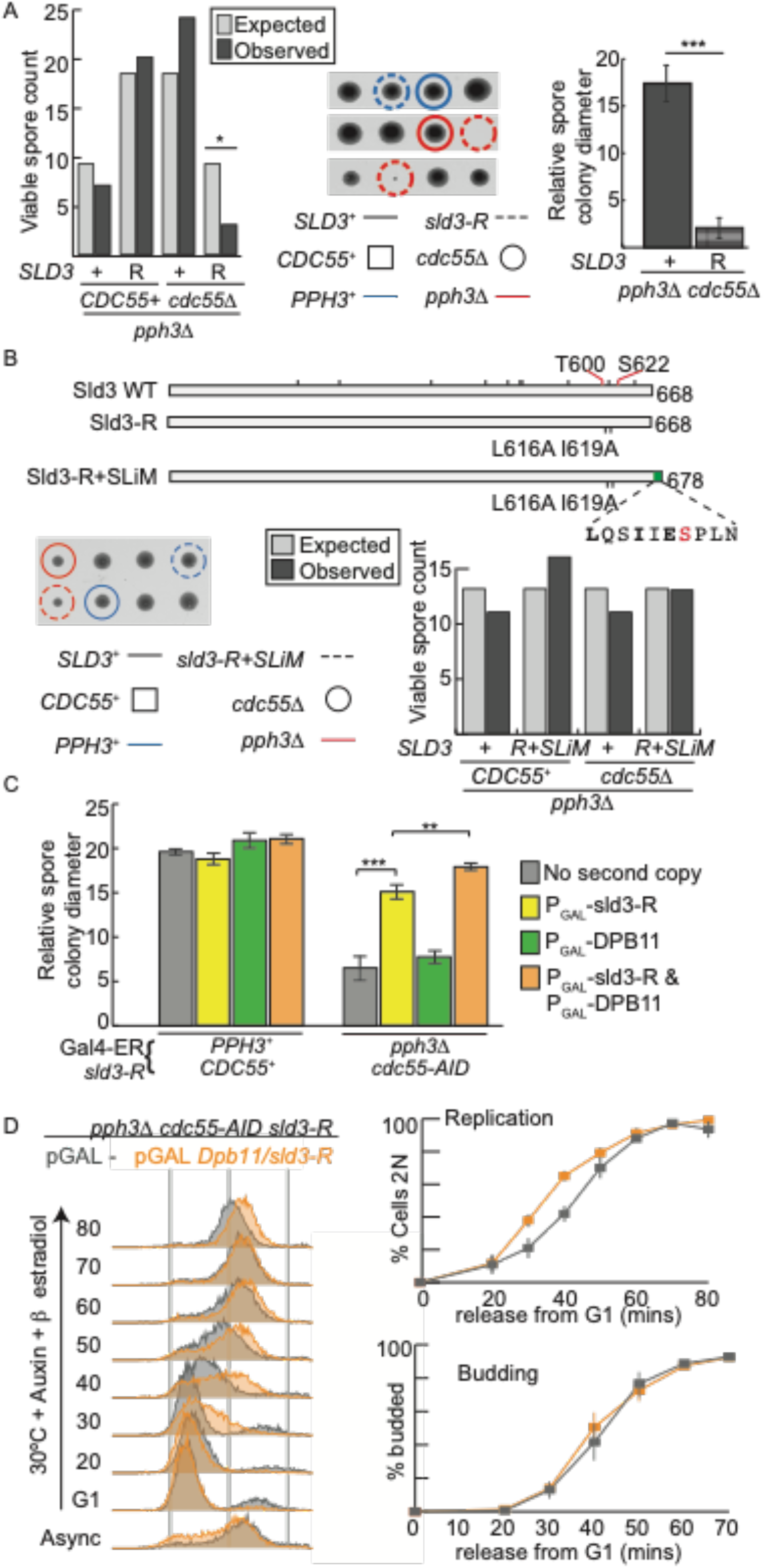
Sld3/Sld2 dephosphorylation is essential. A. Left: Analysis of viable spore count of the indicated strains (observed), compared to the expected number from the Mendelian ratio. All strains are *swe1Δ*. * is a P value of 0.0339 from a Chi-squared test. Middle: example tetrads from the cross on the left. Right: analysis of the relative colony diameter of the indicated genotypes on the tetrad dissection plate (middle). Error bars are SD, *** represents P value <10^−10^ from a t-test. B. Top: Schematic diagram of Sld3. Addition of the Rts1 binding sequence L616-N625 to the very C-terminus of *sld3-R*, generates the *sld3-R+SLiM* allele. Bottom: as A. C. Analysis of relative colony diameter as in A, in the presence of β estradiol + auxin. These strains are all *swe1Δ* and express osTIR1 and Gal4-ER (the P_GAL_ promoter transcription factor as a fusion to ER), allowing the β estradiol induction of P_GAL_. This method was used because *CDC55* mutant strains have defects in growth on galactose. *P_GAL_ sld3-R* and *P_GAL_ DPB11* are expressed as second copies. Error bars are SD, *** and ** represents P values of <0.001 and <0.002 respectively from t-tests. D. Left: Flow cytometry of the indicated strains arrested in G1 at 30°C in the presence of β estradiol + auxin and released into S-phase. These strains are all *swe1Δ* and express osTIR1 and Gal4-ER. Right: quantification of the mean value of the histograms (Replication) and budding from the experiment on the left. Error bars are SD, n=3.

If the phenotypes of *sld3-R* are due to loss of Rts1 binding (Figure 4B), rather than another function of Sld3, then we reasoned that restoring an Rts1 binding site to the *sld3-R* mutant should rescue these phenotypes. Addition of the Rts1 SLiM of Sld3 (L616-N625) to the C-terminus of the *sld3-R* mutant (referred to as *sld3-R+SLiM*, Figure 7B) led to a complete rescue of spore viability and colony size when combined with *cdc55Δ pph3Δ* (Figure 7B and Supplementary Figure 7A). This demonstrates that the lethality of the *sld3-R* mutant in combination with loss of PP2A^CDC55^/PPH3 is very likely due to loss of a direct interaction between Sld3 with Rts1.

Since defects in the dephosphorylation of Sld2 and Sld3 leads to excess CDK phosphorylation of these targets, we wondered whether the partial inhibition of CDK might also counterbalance the defects in dephosphorylation (Supplementary Figure 7B). The *cdc28-as1* mutant has reduced activity even in the absence of 1-NM-PP1 ^9^ and importantly we observed that this allele partially suppressed the growth defect of the *cdc55Δ pph3Δ sld3-R* mutations (Supplementary Figure 7C). This suppression by *cdc28-as1* explains how we obtained viable *cdc55Δ pph3Δ sld3-R* mutants for the analysis of dephosphorylation (e.g in Figure 3/4) and also strongly suggests that it is the hyper-phosphorylation of Sld3/Sld2 by CDK that causes the synthetic lethality of the combined phosphatase mutants *in vivo*.

As failure to dephosphorylate the pre-IC results in retention of this complex at origins (Figure 6), we hypothesised that over-expression of subsets of pre-IC proteins might interfere with complex stability at origins, which might lead to suppression of the phosphatase mutant phenotype. To test this, we over-expressed the Sld3-R mutant that cannot bind to PP2A^RTS1^ with and without over-expression of Dpb11. Over-expression of Sld3-R significantly rescued the growth defect associated with the *cdc55-AID pph3Δ sld3-R* strain, which was further enhanced by over-expression of Dpb11 (Figure 7C). This over-expression also rescued the S-phase defect associated with failure to dephosphorylate Sld3 and Sld2, without affecting the G1-S transition (Figure 7D). Together this is consistent with pre-IC complex turnover at origins being a critical function for Sld3/Sld2 dephosphorylation *in vivo*.

## Discussion

### Dephosphorylation of Sld2/Sld3 during S-phase

Across eukaryotes, CDK is essential for replication initiation and in budding yeast this is mediated by phosphorylation of Sld2 and Sld3. Here we show that these substrates are also actively and specifically dephosphorylated during S-phase, suggesting that the flux of phosphorylation-dephosphorylation is an important feature of these substrates ^32^. The importance of this control is underlined by the fact that three separate phosphatases, PP2A^RTS1^, PP2A^CDC55^ and PP4 (Pph3) are responsible for the dephosphorylation of Sld3 and at least partially of Sld2. One important function for rapid dephosphorylation of Sld2/Sld3 is to allow the release of the pre-IC for efficient origin firing genome-wide (Figure 5/6). This positive role for phosphatases in replication initiation contrasts with the conventional view that phosphatases merely oppose kinase function. Further biochemical studies will be required to determine the exact mechanics of how Sld3/Sld2 dephosphorylation is regulated, how it causes dissociation of the pre-IC and why this dissociation is important for origin firing, but it is intriguing that the Rts1 binding site on Sld3 overlaps with the critical Dpb11 binding site (Figure 4A), suggesting that phosphatase regulation and pre-IC formation are indeed mutually exclusive.

Although Sld3 and Sld2 are retained at early origins in the *cdc55-AID pph3-AID sld3-R* strain, they are still released over time (Figure 6B/C). This might be due to the penetrance of our alleles, in particular due to the still considerable dephosphorylation of Sld2 in this background (Figure 4F), and it is also possible that even in the absence of dephosphorylation there is still some off-rate of the pre-IC from origins. This latter possibility would explain why initiation does occur at a single origin in the reconstituted replication system *in vitro*, in the absence of phosphatase activity ^33^, but *in vivo* these phosphatases are clearly very important for origin firing genome-wide (Figure 5).

### The dual role of dephosphorylation of Sld3/Sld2 throughout the cell cycle

Licensing factors, such as Orc6, must be phosphorylated and inhibited by CDK before Sld3 and Sld2 become phosphorylated in order to ensure that origins only fire once in S-phase ^1^. PP2A has been shown to contribute to the ordering of dephosphorylation of CDK substrates during mitotic exit ^34, 35^ and during interphase ^24, 36^. Loss of PP2A/PP4 resulted in Sld3 hyperphosphorylation in G1-phase (Figure 4D), strongly suggesting that these phosphatases are critical to dephosphorylate Sld3 during mitotic exit. We also show that when CDK activity is interrupted in interphase, for example in a *clb5Δ* strain, the rapid and specific dephosphorylation of Sld3/Sld2 by PP2A/PP4 prevents re-replication by ensuring that these initiation factors are inactivated, while other CDK targets such as Orc6 remain phosphorylated (Figure 2A). The presence of this Sld2/3 specific phosphatase activity explains the continual requirement for CDK activity for origin firing throughout S-phase ^14^ and may explain how multiple organisms, from fission yeast to humans, respond to DNA damage in S-phase by inhibiting CDK, without causing subsequent re-replication ^37, 38^. This study therefore highlights a dual function for PP2A/PP4 phosphatases in replication control: to allow pre-IC recycling and origin firing in S-phase (Figure 5/6), and to ensure that Sld2/Sld3 are rapidly dephosphorylated to prevent inappropriate initiation at any time when CDK activity drops, for example during mitotic exit (Figure 4D).

Although PP2A/PP4 are required for Sld3 dephosphorylation in M/G1 phase, we believe that it is the replication initiation defects in S-phase (Figure 5/6) that cause the loss of viability in the *cdc55Δ pph3Δ sld3-R* strain (Figure 7). Firstly, we see no evidence for re-replication in G1 phase in this strain, as determined by flow cytometry and DNA copy number analysis (data not shown), probably because although Sld3 remains phosphorylated in G1 phase in this strain, Sld2 is not (Figure 4D and 4F). In addition, we can suppress both the growth defect and S-phase defect by over-expression of Sld3/Dpb11 (Figure 7D), which may help to drive the dissolution of the pre-IC at origins.

### Conservation of PP2A function in replication control

It has been shown in Xenopus egg extracts that PP2A plays a positive role in replication initiation ^36, 39^, critically during the CDK-dependent step of Cdc45 recruitment to chromatin ^40^. It remains to be tested whether PP2A in vertebrates regulates pre-IC dynamics through dephosphorylation of the essential CDK targets. Analysing this in human cells may be complicated by a potential negative role for PP2A in origin firing through the degradation of the Sld3 orthologue Treslin ^41^, which we do not observe in yeast (Supplementary Figure 5F). Given the importance of inhibition of DNA replication as a chemotherapeutic strategy, the novel essential function for PP2A/PP4 in replication initiation described here may provide a new rationale for targeting these phosphatases in cancers ^42^.

## Methods

### Cell synchrony experiments, growth assays and yeast strains

The yeast strains used in this work are listed in Supplementary Table 1. Unless otherwise stated, yeast cells were grown at 25°C in YPAD (1% yeast extract, 2% peptone, 0.004% adenine sulfate, 2% glucose) medium. For cell synchronisation experiments, overnight cultures of MATa strains were grown to 1.0 × 10^7^ cells/ml in YPAD at 25°C. Cultures were blocked by addition of 5 μg/ml of α factor (GenScript) every 90 min in YPAD until <10% cells were budded. Cells were released from the arrest by washing and then placed in fresh YPAD medium. For G2 arrest, Nocodazole was used at a concentration of 10 μg/ml in dimethyl sulfoxide (DMSO) and re-added after two hours where necessary. Where appropriate: *cdc28-as1* was inhibited by adding 5μM 1-NMPP1 (Merck); auxin-inducible degrons were induced with 1mM indole-3-acetic acid (IAA, Sigma-Aldrich) for 30 minutes at 37°C; the DNA damage checkpoint was activated by release of cells into YPAD medium containing 200mM hydroxyurea (HU, Acros Organics).

### Protein extraction, western blotting and phosphorylation analysis

Protein was extracted from pelleted yeast cells by bead beating with glass beads in 20% TCA. The resulting precipitate was pelleted and resuspended in Laemmli buffer, neutralised with Tris base and boiled. For Rad53, Orc6 and Mcm4 phosphorylation, 7.5%, 10% and 6% SDS PAGE gels were used respectively. For higher resolution of protein phosphorylation, Phos-tag (Alpha Laboratories) SDS PAGE was employed. Orc6 (for Figure 1C) was resolved on 8% SDS-PAGE with 20 μM Phos-tag; Sli15 and Yen1 were separated on a 4% SDS-PAGE gel with 12.5 μM Phos-tag and for Sld2 and Sld3, a 4% SDS-PAGE gel with 25 μM Phos-tag was used. For Western blotting, the Phos-tag gels were incubated 4 × 10 min in 100 mM EDTA before electrophoretic transfer onto Nitrocellulose membrane (Sigma). Proteins were detected by immunoblotting using the following primary antibody concentrations in 5% milk (VWR)/Tris Buffered Saline -Tween20 (Sigma) : anti-Myc 1:10,000 (9E10, Roche, Cat no: 11 667 149 001; RRID: <u>AB_390912</u>) or (Abcam, Cat no: ab9132; RRID: AB_307033); anti-HA 1:1000 (16B12, AbCam); anti-Orc6 1:2000 (Stillman lab); anti-Rad53 1:5000 (ab104232, Abcam, Cat no: ab104232; RRID: <u>AB_2687603</u>); anti-Sld2 1:1000 (Zegerman lab); anti-Clb2 1:5000 (Santa Cruz); anti-flag 1:1000 (Sigma, cat no: F3165; RRID: RRID:AB_259529); anti-Mcm4 1:2000 (cat no: sc-166037, RRID:AB_2012315); anti-GST (Abcam, cat. no: ab58626, RRID: AB_880249). Blots were visualised using ECL or ECL prime system (Amersham) and quantified using FIJI.

### Flow cytometry, Budding Index and T_rep_ analysis

For flow cytometry, 500μl of culture was pelleted and fixed in 70% ethanol overnight at 4°C. The pellet was washed once with 0.05M Tris pH8.0 and re-suspended in RNAse A (10μg/ml in 0.05M Tris pH8.0, Sigma-Aldrich) for overnight 37°C incubation. At this point samples were taken to count population Budding Index. The remaining sample was re-suspended in pepsin (5mg/ml in 50mM HCl, Sigma-Aldrich) for 30 minutes at 37°C and washed once again with 0.05M Tris pH8.0. DNA staining was then performed re-suspension in propidium iodide (0.5μg/ml in 0.05M Tris pH8.0, Sigma-Aldrich). Diluted samples were sonicated and 10,000 cells were analysed on a FACS Calibur (BD). Flow cytometry profiles were then visualised using FlowJo v10.7.1 (FlowJo, LLC).

For T_rep_ profile generation, yeast genomic DNA was extracted from 5 minute interval samples using the smash and grab method (https://fangman-brewer.genetics.washington.edu/smash-n-grab.html). DNA was sonicated (Bioruptor Pico, Diagenode), and the libraries were prepared according to the TruSeq Nano sample preparation guide from Illumina. To generate replication timing profiles, the ratio of uniquely mapped reads in the replicating samples to the non-replicating samples was calculated following ^43^. T_rep_ was determined as the time-point (in minutes after G1 release) at which each genomic bin is halfway from one copy to two copies ^44^. Replication profiles were generated using ggplot and smoothed using a moving average in R. The values of T_rep_ for Fig. 5F were taken from OriDB ^45^.

### CHIP-seq, CHIP-qPCR

For Mcm4 and Sld3 CHIP, 200 mL of yeast culture at 1×10^7^ cells/ml was crosslinked with formaldehyde (final concentration 1%) for 25 minutes at room temperature with gentle rotation. For Sld2 CHIP, crosslinking was performed with 1.5mM EGS (ethylene glycol bis(succinimidyl succinate)) for 10 minutes, followed by 1% Formaldehyde for 10 minutes. Crosslinking reactions were terminated by addition of 125 mM Glycine for 5 minutes at room temperature with gentle rotation. Cells were washed once with PBS, once with 50 mM HEPES and resuspended in lysis buffer (50mM HEPES/KOH pH7.5, 1mM EDTA, 1% Triton X-100, 0.1% Sodium deoxycholate, 140mM NaCl, Protease Inhibitor Cocktail (Sigma), Protease inhibitors (Roche)). 300μL of glass beads were added and cells were mechanically disrupted with tissue homogenizer at 4°C (Precellys) for 6 cycles at 6000 rpm for 30 s, with 3 minutes incubation on ice between each cycle. Cell lysates were sonicated 18 cycles of 30 s on and 30 s off (Bioruptor Pico, Diagenode) and insoluble material was discarded by centrifugation at 13000rpm for 20 minutes at 4°C. Supernatants were transferred to a new tube and spiked with lysate of a strain containing epitope-tagged Cse4 for normalisation purposes. Samples were then added to pre-blocked antibody-conjugated magnetic beads (anti-HA Magnetic beads, Pierce cat no: 88836; or anti-Myc PL14, MBL international Cat no: M047-3S RRID: AB_591108, with Protein A dynabeads, Thermofisher) and were incubated with rotation overnight at 4°C. Beads were collected and washed once with lysis buffer, once with buffer 1 (50mM HEPES/KOH pH7.5, 1mM EDTA, 1% Triton X-100, 0.1% Sodium deoxycholate and 250mM NaCl), once with buffer 2 (50mM HEPES/KOH pH7.5, 1mM EDTA, 1% Triton X-100, 0.1% Sodium deoxycholate and 500mM NaCl), once with buffer 3 (0.25M LiCl, 0.5% NP-40, 0.5% Sodium deoxycholate, 1mM EDTA, 10mM Tris-HCl pH8) and once with TE pH8. Samples were eluted in elution buffer (0.85X TE pH8, 1% SDS, 0.25M NaCl) for 30 minutes at 65°C. Eluted materials were transferred to new tubes and treated with RNase A (100 μg/ml; Roche) for 2.5 hours at 37°C, then with Proteinase K (800 μg/ml; Roche) overnight at 65°C. DNA was purified with CHIP clean concentrator kit (Zymo). For CHIP-qPCR: qPCR was performed on diluted samples using LightCycler 480 SYBR green 1 master kit (Roche) using primers listed below.

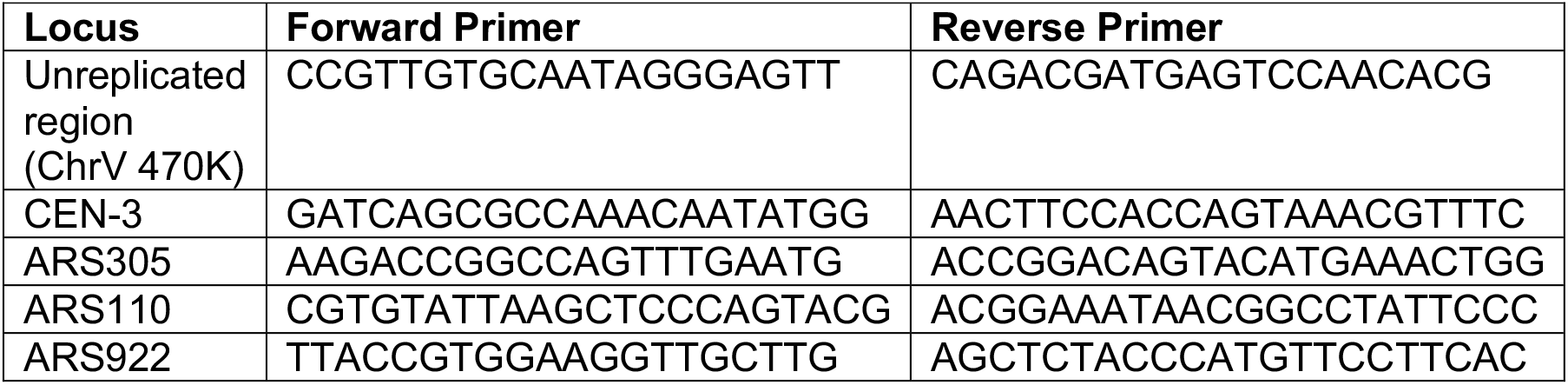

Using the 2^ΔΔct^ method, CHIP recovery relative to Input was normalised to the corresponding recovery from an unreplicated region and/or centrometric region to calculate fold enrichment per sample. For each strain, enrichment was then reported relative to the G1 sample.

For CHIP-seq: Sequencing libraries were prepared according to instructions of Illumina Truseq-Nano except: size selection was adjusted for 200bp fragments; adaptors were used at 1:100 dilution and 14-16 cycles of amplification were performed. Sequencing was performed using an Illumina Hiseq 1500 or NovaSeq6000. Two biological replicates were performed for all experiments.

CHIP-seq data was analysed as follows: Sequence reads were mapped using Bowtie2 (version 2.2.6). Alignment files (BAM and SAM) were sorted and indexed using samtools (version 0.1.19-96b5f2294a). CHIP and corresponding input BAM files were compared using bamCompare (version 3.0.2). To compare the BAM files, the genome is partitioned into bins of equal size, then the number of reads found in each bin is counted per file, and finally the log2 ratio is reported. The CHIP and input are normalised for sequencing depth. Heatmaps were generated using computeMatrix and plotHeatmap (version 3.0.2).

### Peptide pulldown experiments

pET21b Dpb11(1-395) GST ^11^ was transformed into E. coli cells (FB850). The cells were cultured at 37°C in LB medium (1% peptone, 0.5% yeast extract and 0.5% NaCl), with 25μg/ml chloramphenicol and 100μg/ml ampicilin, to an OD600 of 0.5. Protein production was induced by adding 1mM isopropyl-β-D-thiogalactoside (IPTG) and culturing the cells at 16°C overnight. The cells were harvested by centrifugation, resuspended in pre-cooled lysis buffer (20mM HEPES pH 7.4, 200mM NaCl, 5% glycerol, 0.5mM EDTA, 1 mM phenylmethylsulfonyl fluoride (PMSF), Benzamidine HCl, Leupeptin, Pepstatin A) and sonicated 4 × 30s on/off. The extracts were cleared by centrifugation (48,000g, 4°C, 20min), and the supernatant was loaded on a GSTrap column (AKTA, GE HEalthcare) at 4°C following the manufacturer’s instructions. Proteins were eluted using lysis buffer containing 25mM reduced Glutathione. Fractions were collected, checked by SDS-PAGE and Coomassie blue staining, and pure fractions were pooled and aliquoted for storage at −80°C.

For the pulldown, 100pmol biotinylated peptides (Peptide 2.0) were bound to 8μl streptavidin M-280 Dynabeads (Thermo Fisher Scientific) for 30mins in PBS at room temperature with agitation. After two washes with HBS/BSA (10mM HEPES pH 7.4, 150mM NaCl / 5mg/ml BSA), 200μl of 1μM GST-Dpb11 1-395 diluted in HBS + 3mM EDTA was added and beads were incubated for 30 minutes at room temperature. After three washes with 1ml HBS, beads were eluted in 30μl Laemmli buffer by boiling for 10 minutes at 95°C.

## Data and Software Availability

Sequencing data is available at GEO: <u>GSE186490</u>.

## Acknowledgements

We are grateful to Mike Stark, Douglas Kellogg, Adam Rudner and Joao Matos for strains. We thank members of the Zegerman lab for critical reading of the manuscript and Mark Johnson for help with analysis of the replication profiles. Work in the PZ lab was supported by AICR 10-0908, Wellcome Trust 107056/Z/15/Z, Cancer Research UK C15873/A12700 and Gurdon Institute funding (Cancer Research UK C6946/A14492, Wellcome Trust 092096). MS was funded by the BBSRC BB/M011194/1. FJ was funded by AstraZeneca studentship CR-000608. BS was funded by Der Schweizerische Nationalfonds (SNF PBEZP3-135429) and European Molecular Biology Organisation (EMBO ALT 646-2011).

## Author Contribution

All authors performed and designed the experiments. PZ wrote the paper.

## Declaration of Interests

The authors declare no conflicts of interest.

## Notes

https://www.ncbi.nlm.nih.gov/geo/query/acc.cgi?acc=GSE186490

### Competing Interest Statement

The authors have declared no competing interest.

## References

1. Siddiqui, K., On, K.F. & Diffley, J.F. Regulating DNA replication in eukarya. Cold Spring Harbor perspectives in biology 5 (2013).

2. Bell, S.P. & Labib, K. Chromosome Duplication in Saccharomyces cerevisiae. Genetics 203, 1027–1067 (2016).

3. Miyazawa-Onami, M., Araki, H. & Tanaka, S. Pre-initiation complex assembly functions as a molecular switch that splits the Mcm2-7 double hexamer. EMBO reports 18, 1752–1761 (2017).

4. Tanaka, S. & Araki, H. Helicase activation and establishment of replication forks at chromosomal origins of replication. Cold Spring Harbor perspectives in biology 5, a010371 (2013).

5. Boos, D. & Ferreira, P. Origin Firing Regulations to Control Genome Replication Timing. Genes 10 (2019).

6. Mantiero, D., Mackenzie, A., Donaldson, A. & Zegerman, P. Limiting replication initiation factors execute the temporal programme of origin firing in budding yeast. The EMBO journal 30, 4805–4814 (2011).

7. Tanaka, S., Nakato, R., Katou, Y., Shirahige, K. & Araki, H. Origin association of Sld3, Sld7, and Cdc45 proteins is a key step for determination of origin-firing timing. Current biology : CB 21, 2055–2063 (2011).

8. Lynch, K.L., Alvino, G.M., Kwan, E.X., Brewer, B.J. & Raghuraman, M.K. The effects of manipulating levels of replication initiation factors on origin firing efficiency in yeast. PLoS genetics 15, e1008430 (2019).

9. Bishop, A.C. et al. A chemical switch for inhibitor-sensitive alleles of any protein kinase. Nature 407, 395–401 (2000).

10. Tanaka, S. et al. CDK-dependent phosphorylation of Sld2 and Sld3 initiates DNA replication in budding yeast. Nature 445, 328–332 (2007).

11. Zegerman, P. & Diffley, J.F. Phosphorylation of Sld2 and Sld3 by cyclin-dependent kinases promotes DNA replication in budding yeast. Nature 445, 281–285 (2007).

12. Tak, Y.S., Tanaka, Y., Endo, S., Kamimura, Y. & Araki, H. A CDK-catalysed regulatory phosphorylation for formation of the DNA replication complex Sld2-Dpb11. The EMBO journal 25, 1987–1996 (2006).

13. Jackson, L.P., Reed, S.I. & Haase, S.B. Distinct mechanisms control the stability of the related S-phase cyclins Clb5 and Clb6. Molecular and cellular biology 26, 2456–2466 (2006).

14. Donaldson, A.D. et al. CLB5-dependent activation of late replication origins in S. cerevisiae. Mol Cell 2, 173–182 (1998).

15. Zhai, Y., Yung, P.Y., Huo, L. & Liang, C. Cdc14p resets the competency of replication licensing by dephosphorylating multiple initiation proteins during mitotic exit in budding yeast. Journal of cell science 123, 3933–3943 (2010).

16. Jin, F. et al. Temporal control of the dephosphorylation of Cdk substrates by mitotic exit pathways in budding yeast. Proceedings of the National Academy of Sciences of the United States of America 105, 16177–16182 (2008).

17. Stegmeier, F. & Amon, A. Closing mitosis: the functions of the Cdc14 phosphatase and its regulation. Annual review of genetics 38, 203–232 (2004).

18. Akiyoshi, B. & Biggins, S. Cdc14-dependent dephosphorylation of a kinetochore protein prior to anaphase in Saccharomyces cerevisiae. Genetics 186, 1487–1491 (2010).

19. Zegerman, P. & Diffley, J.F. Checkpoint-dependent inhibition of DNA replication initiation by Sld3 and Dbf4 phosphorylation. Nature 467, 474–478 (2010).

20. Lopez-Mosqueda, J. et al. Damage-induced phosphorylation of Sld3 is important to block late origin firing. Nature 467, 479–483 (2010).

21. Rodriguez, C.E., Sobol, Z. & Schiestl, R.H. 9,10-Phenanthrenequinone induces DNA deletions and forward mutations via oxidative mechanisms in the yeast Saccharomyces cerevisiae. Toxicology in vitro : an international journal published in association with BIBRA 22, 296–300 (2008).

22. Kalev, P. & Sablina, A.A. Protein phosphatase 2A as a potential target for anticancer therapy. Anti-cancer agents in medicinal chemistry 11, 38–46 (2011).

23. Yang, H., Jiang, W., Gentry, M. & Hallberg, R.L. Loss of a protein phosphatase 2A regulatory subunit (Cdc55p) elicits improper regulation of Swe1p degradation. Molecular and cellular biology 20, 8143–8156 (2000).

24. Godfrey, M. et al. PP2A(Cdc55) Phosphatase Imposes Ordered Cell-Cycle Phosphorylation by Opposing Threonine Phosphorylation. Mol Cell 65, 393–402 e393 (2017).

25. Moyano-Rodriguez, Y. & Queralt, E. PP2A Functions during Mitosis and Cytokinesis in Yeasts. International journal of molecular sciences 21 (2019).

26. Ronne, H., Carlberg, M., Hu, G.Z. & Nehlin, J.O. Protein phosphatase 2A in Saccharomyces cerevisiae: effects on cell growth and bud morphogenesis. Molecular and cellular biology 11, 4876–4884 (1991).

27. Evans, D.R. & Stark, M.J. Mutations in the Saccharomyces cerevisiae type 2A protein phosphatase catalytic subunit reveal roles in cell wall integrity, actin cytoskeleton organization and mitosis. Genetics 145, 227–241 (1997).

28. Wang, X., Bajaj, R., Bollen, M., Peti, W. & Page, R. Expanding the PP2A Interactome by Defining a B56-Specific SLiM. Structure 24, 2174–2181 (2016).

29. Hertz, E.P.T. et al. A Conserved Motif Provides Binding Specificity to the PP2A-B56 Phosphatase. Mol Cell 63, 686–695 (2016).

30. Kennedy, E.K. et al. Redundant Regulation of Cdk1 Tyrosine Dephosphorylation in Saccharomyces cerevisiae. Genetics 202, 903–910 (2016).

31. Wei, H. et al. Carboxymethylation of the PP2A catalytic subunit in Saccharomyces cerevisiae is required for efficient interaction with the B-type subunits Cdc55p and Rts1p. The Journal of biological chemistry 276, 1570–1577 (2001).

32. Gelens, L. & Saurin, A.T. Exploring the Function of Dynamic Phosphorylation-Dephosphorylation Cycles. Dev Cell 44, 659–663 (2018).

33. Yeeles, J.T., Deegan, T.D., Janska, A., Early, A. & Diffley, J.F. Regulated eukaryotic DNA replication origin firing with purified proteins. Nature 519, 431–435 (2015).

34. Touati, S.A. et al. Cdc14 and PP2A Phosphatases Cooperate to Shape Phosphoproteome Dynamics during Mitotic Exit. Cell reports 29, 2105–2119 e2104 (2019).

35. Holder, J., Poser, E. & Barr, F.A. Getting out of mitosis: spatial and temporal control of mitotic exit and cytokinesis by PP1 and PP2A. FEBS letters 593, 2908–2924 (2019).

36. Krasinska, L. et al. Protein phosphatase 2A controls the order and dynamics of cell-cycle transitions. Mol Cell 44, 437–450 (2011).

37. Kumar, S. & Huberman, J.A. Checkpoint-dependent regulation of origin firing and replication fork movement in response to DNA damage in fission yeast. Molecular and cellular biology 29, 602–611 (2009).

38. Bartek, J., Lukas, C. & Lukas, J. Checking on DNA damage in S phase. Nature reviews. Molecular cell biology 5, 792–804 (2004).

39. Lin, X.H. et al. Protein phosphatase 2A is required for the initiation of chromosomal DNA replication. Proceedings of the National Academy of Sciences of the United States of America 95, 14693–14698 (1998).

40. Chou, D.M., Petersen, P., Walter, J.C. & Walter, G. Protein phosphatase 2A regulates binding of Cdc45 to the prereplication complex. The Journal of biological chemistry 277, 40520–40527 (2002).

41. Charrasse, S. et al. Ensa controls S-phase length by modulating Treslin levels. Nature communications 8, 206 (2017).

42. O’Connor, C.M., Perl, A., Leonard, D., Sangodkar, J. & Narla, G. Therapeutic targeting of PP2A. The international journal of biochemistry & cell biology 96, 182–193 (2018).

43. Batrakou, D.G., Muller, C.A., Wilson, R.H.C. & Nieduszynski, C.A. DNA copy-number measurement of genome replication dynamics by high-throughput sequencing: the sort-seq, sync-seq and MFA-seq family. Nat Protoc 15, 1255–1284 (2020).

44. Raghuraman, M.K. et al. Replication dynamics of the yeast genome. Science 294, 115–121 (2001).

45. Siow, C.C., Nieduszynska, S.R., Muller, C.A. & Nieduszynski, C.A. OriDB, the DNA replication origin database updated and extended. Nucleic Acids Res 40, D682–686 (2012).

