## Supplementary material for "Dephosphorylation of the pre-initiation complex during S-phase is critical for origin firing": Supplemetal Figures

Supplementary Figure 1

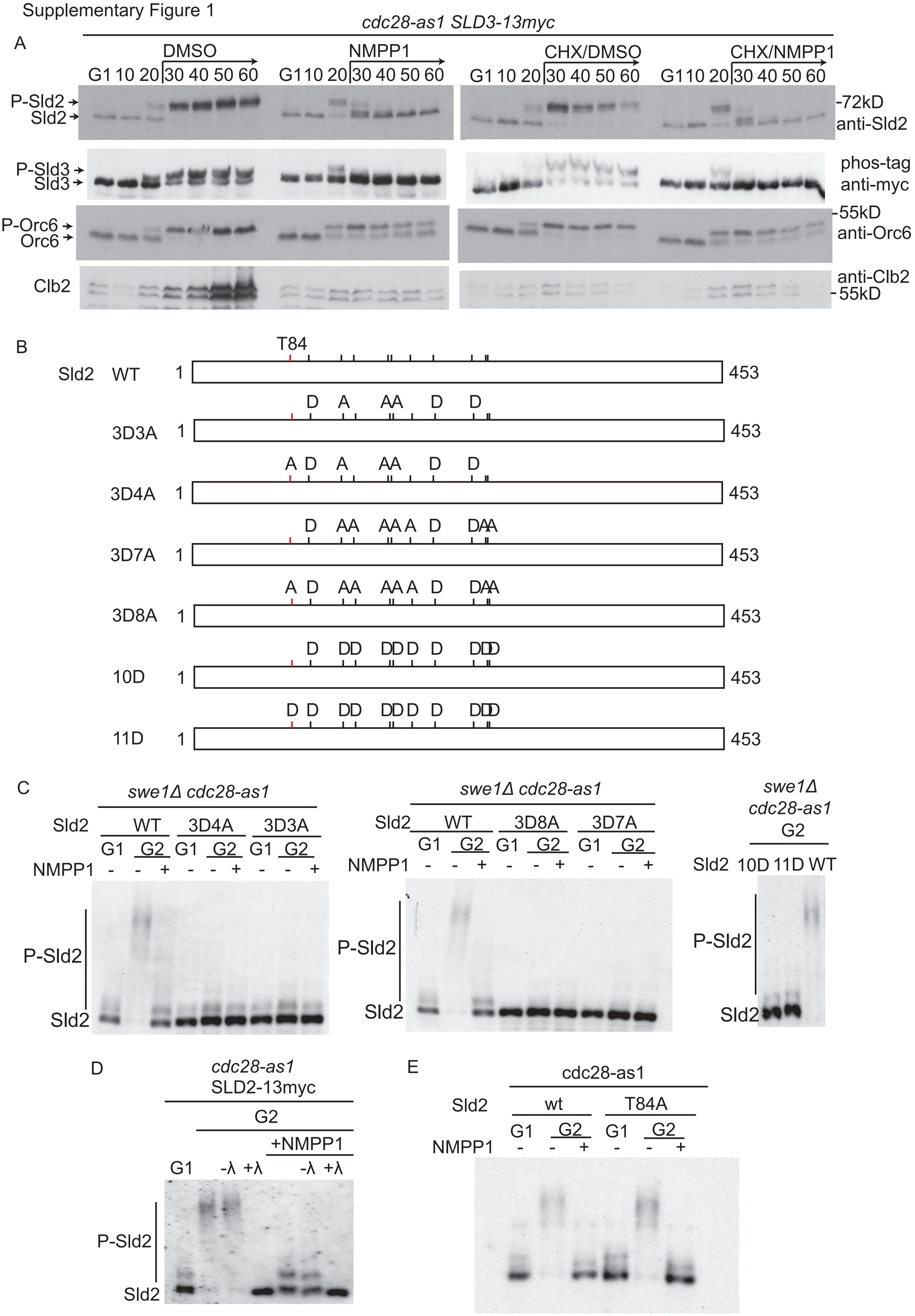

**Supplementary Figure 1.**

A. Western blots of the indicated strains arrested in G1 phase with alpha factor and released into S-phase for the indicated time. The *cdc28-as1* inhibitor 1-NM-PP1 and/or cycloheximide (CHX) was added at 25 minutes. 1-NM-PP1 is dissolved in DMSO, so this solvent is added as a control. The translation of the mitotic cyclin Clb2 demonstrates the efficacy of cycloheximide.
B. Top: scale diagram of Sld2, with the 11 CDK sites indicated by lines. The essential CDK site T84 is indicated in red.
C. Anti-myc phos-tag western blots of strains containing the indicated Sld2-13myc alleles described in B as a second copy. Cells were either arrested in G1 phase or in G2 phase with nocodazole, with (+) or without (-) the addition of 1-NM-PP1 for 10 minutes. Note that all alleles, except for wild type showed a complete lack of CDK phosphorylation in G2, similar to the 11D mutant (right) which lacks all CDK sites.
D. Inhibition of CDK, leads to rapid dephosphorylation of Sld2 (just as it does for Sld3), but the dephosphorylated Sld2 runs as a doublet (see C above, but also main figures including Figure 3F). To address the nature of this doublet we added $\lambda$ phosphatase to the G2 extracts from C, before and after addition of 1-NM-PP1. Interestingly the doublet disappears in the presence of $\lambda$ phosphatase, suggesting Sld2 retains a phosphorylated site even after the addition of 1-NM-PP1. This site is unlikely to be a CDK site, because the 11D mutant (which lacks any CDK sites) and the Sld2 protein in G1 phase also runs as a doublet (C, above). Therefore, Sld2 is phosphorylated by a kinase other than CDK and this phosphorylation is resistant to the rapid dephosphorylation of Sld2 we observe in interphase.
E. As C, but with just the T84A mutant which lacks the essential CDK site.

Supplementary Figure 2

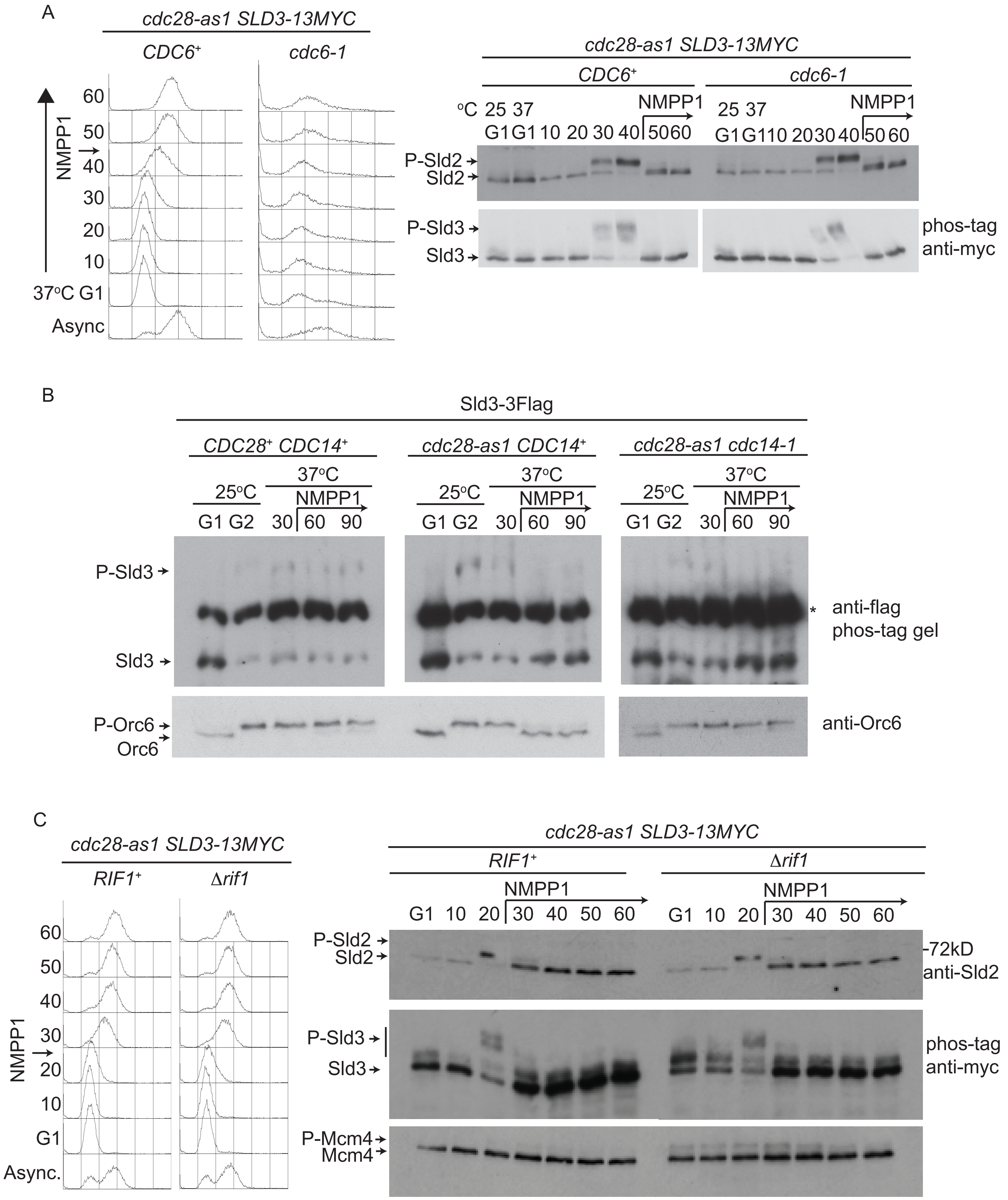

**Supplementary Figure 2.**

A. Flow cytometry (left) and western blots (right) of the indicated strains arrested in G1 phase at 25°C with alpha factor, held in G1 phase for 60 minutes at 37°C and then released into S-phase at 37°C for the indicated time. The *cdc28-as1* inhibitor 1-NM-PP1 was added at 45 minutes.

B. The indicated strains were arrested in G2 phase with nocodazole at 25°C and held in nocodazole while shifted to 37°C for the indicated times. Unfortunately, this anti-flag western blot recognises a non-specific band * immediately between the unphosphorylated and the phosphorylated Sld3. The important point is that the addition of 1-NM-PP1 to the *cdc28-as1* strain either with or without *cdc14-1* results in dephosphorylation of Sld3 (perhaps easiest to observe the accumulation of the unphosphorylated Sld3 band), whilst Orc6 is only dephosphorylated in the *CDC14*$^{+}$ strain.

C. Flow cytometry (left) and western blots (right) of the indicated strains arrested in G1 phase with alpha factor and released into S-phase for the indicated time. The *cdc28-as1* inhibitor 1-NM-PP1 was added at 25 minutes. Note that Mcm4, but also Sld3 are hyper-phosphorylated in G1 phase in the *rif1*Δ strain, both have which have been shown to be DDK-dependent [1]. Upon addition of 1-NM-PP1 both Sld2 and Sld3 are dephosphorylated (including the extra DDK phosphorylation on Sld3 in the *rif1*Δ strain), but Mcm4 is not. This demonstrates that the Sld3/Sld2 phosphatase is specific for Sld3/Sld2 over Mcm4 and is not Rif1-dependent.

Supplementary Figure 3

A Asyn. ⟶ α-factor G1 ⟶ 30 min S ⟶ 10 min with phosphatase inhibitor +NMPP1

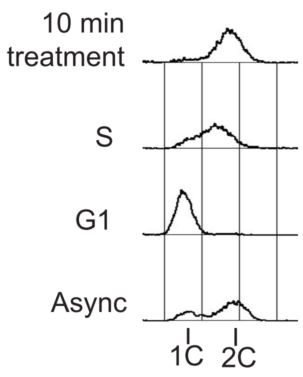


| No. | Phosphatase inhibitor | Target |
|---|---|---|
| 1 | Cantharidin | PP1 and PP2A |
| 2 | FK506 | Calcineurin |
| 3 | Microcystin-LR | PP1 and PP2A |
| 4 | 9, 10 Phenanthrenequinone | Tyrosine phosphatase |
| 5 | Sodium orthovanadate | Alkaline phosphatase |
| 6 | Pentamidine | Tyrosine phosphatase |

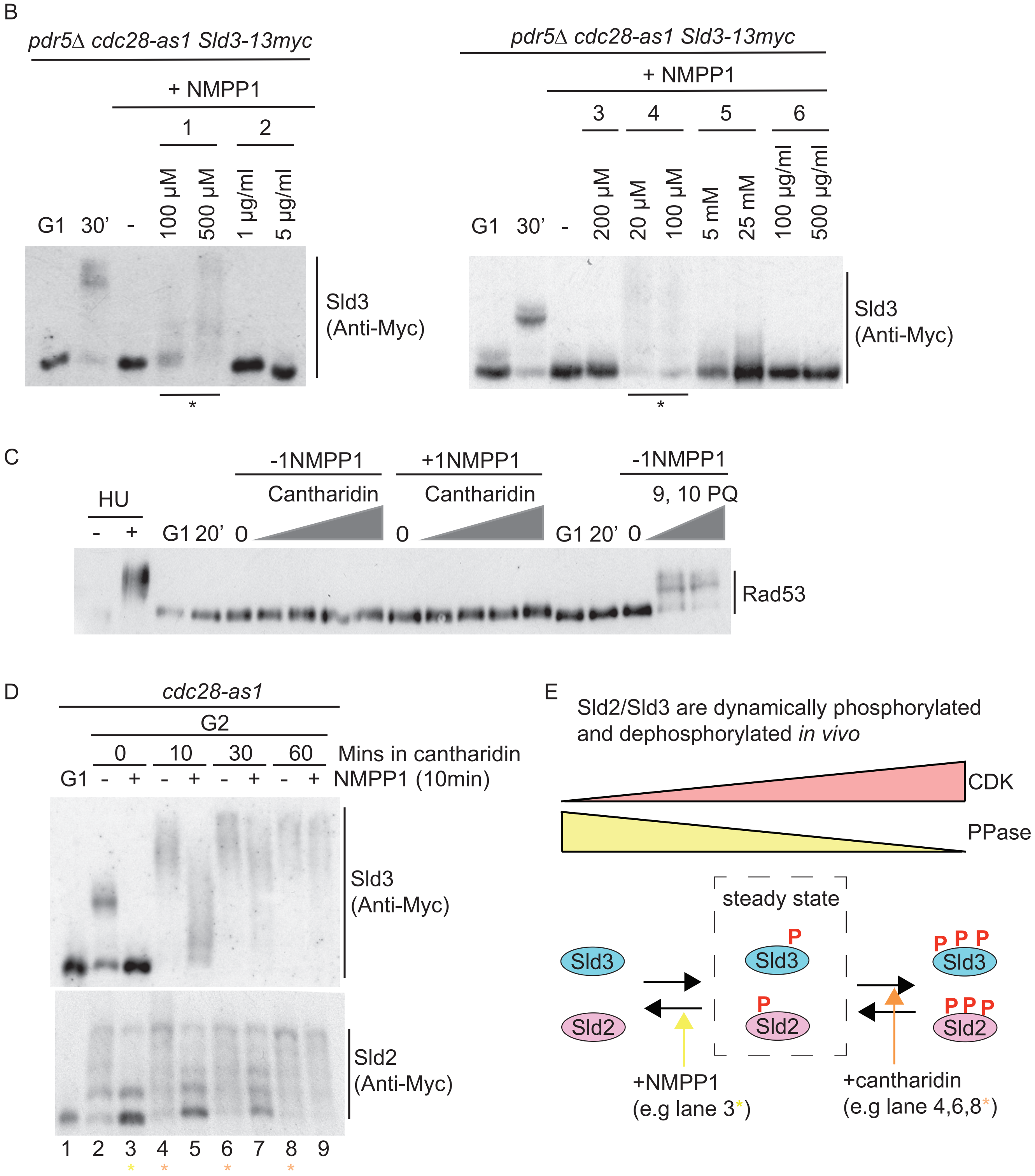

**Supplementary Figure 3.**

A. Top: Schematic diagram of the chemical inhibitor strategy. The *pdr5Δ Sld3-13myc cdc28-as1* strain was arrested in alpha factor, released into S-phase for 30 minutes and then phosphatase inhibitor and 1-NM-PP1 were simultaneously added for 10 minutes. Strains were *pdr5Δ* to potentially increase uptake of phosphatase inhibitors. Bottom: an example flow cytometry of this experiment and the table of broad-specificity inhibitors (1-6) used in this analysis.
B. Western blot in the presence of the inhibitors numbered 1-6 in A. * indicates lanes where 1-NM-PP1 + inhibitor did not result in dephosphorylation.
C. Rad53 western blot of cells in released from G1 into S-phase for 20 minutes, followed by the addition of increasing concentrations of cantharidin up to 500μM or phenanthrenequinone (PQ) up to 100μM or no inhibitor (0) for 10 minutes.
D. Phos-tag western blot of either *Sld2-13myc* or *Sld3-13myc cdc28-as1* strains arrested in G2 phase with nocodazole, then held in G2 phase in the presence of 1000μM cantharidin for 10, 30 or 60 minutes. For the final 10 minutes of these incubations with cantharidin, cells were treated for 10 minutes with DMSO (-) or with 5μM 1-NM-PP1 (+). This experiment therefore tests whether 1-NM-PP1 addition affects the phosphorylation of Sld2/Sld3 if the phosphatase has already been inhibited.
E. Figure D demonstrates that Sld3 and Sld2 are constantly phosphorylated and dephosphorylated *in vivo*. Addition of 1-NM-PP1 leads to rapid dephosphorylation of Sld2/Sld3 (lane 3, D), demonstrating that CDK is continually required to maintain the steady state phosphorylation. Conversely, addition of cantharidin leads to rapid hyperphosphorylation of Sld2/Sld3 (e.g lane 4, D) suggesting that these proteins are continually dephosphorylated *in*

*vivo*. The Sld3/Sld2 hyperphosphorylation caused by cantharidin addition is not specific to the *cdc28-as1* mutant strain, as we observe the same result in *CDC28*$^{+}$ strains (data not shown). The constant flux of CDK and cantharidin-sensitive phosphatase activity towards Sld2/Sld3 leads to a steady state level of phosphorylation of Sld2/Sld3 in interphase.

Supplementary Figure 4

A

Cantharidin $IC_{50}$ (from Kalev and Sablina 2011)

PP2A (0.2nM) < PP1 (0.5-2.0nM) < PP4 (50nM) < PP5 (200nM)

PP2A family

YDL047w SIT4/PPH1 (essential)

YDL134c PPH21 | paralogues (80% sequence identity)

YDL188c PPH22 |

YDR075w PPH3

YNR032w PPG1

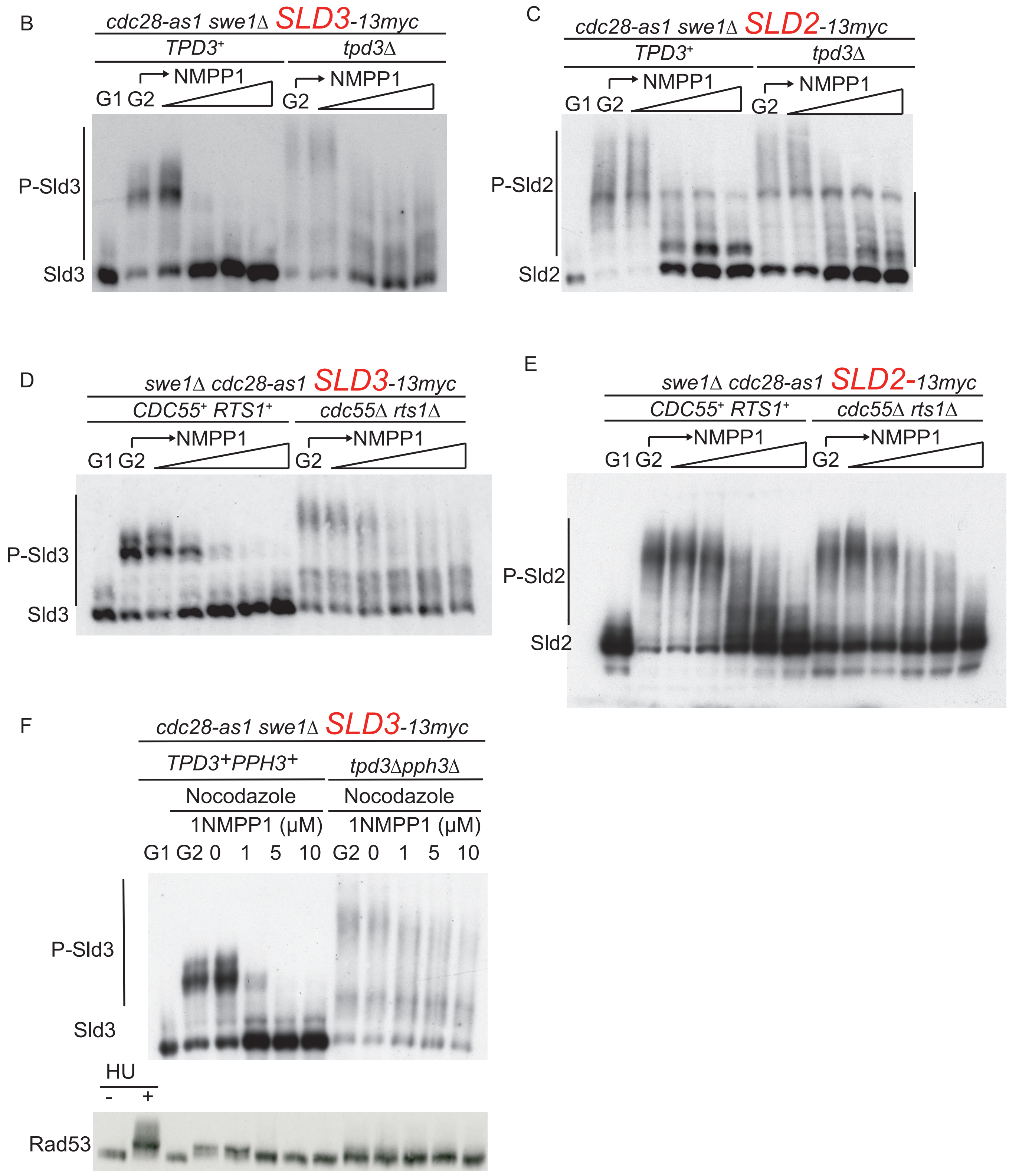

**Supplementary Figure 4.**

A. Left: IC50 values for cantharidin and different phosphatase families from [2]. Right: table of PP2A family phosphatases in budding yeast. Only *SIT4* is an essential gene.

B-F. The indicated strains were arrested in nocodazole (G2) and treated with increasing concentrations of 1-NM-PP1 up to 10μM for 10 minutes for B,C and F and up to 5μM for 10 minutes for D,E.

Supplementary Figure 5

A

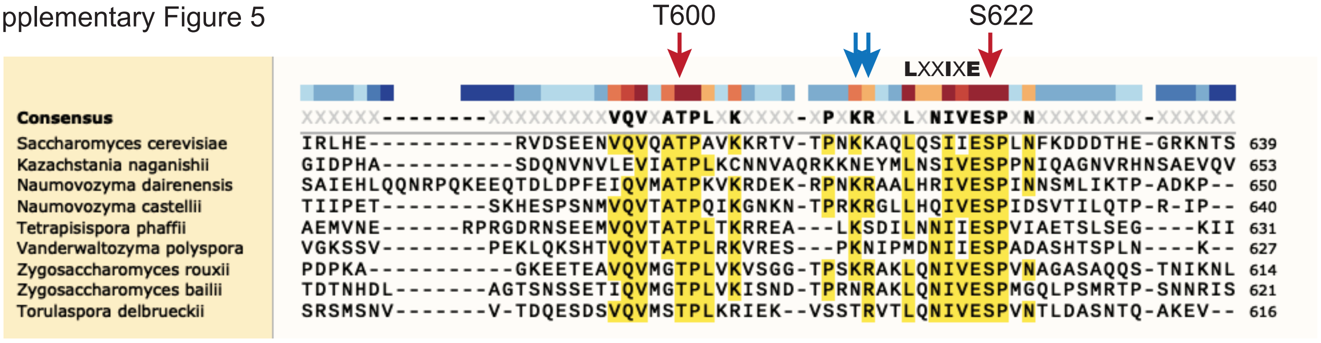


B Sld3 peptides 590-639

1. DSEENVQVQATPAVKKRTVTPNKKAQLQSIIESPLNFKDDDTHEGRKNTS

2. DSEENVQVQA(P)TPAVKKRTVTPNKKAQLQSIIE(P)SPLNFKDDDTHEGRKNTS

3. DSEENVQVQATPAVKKRTVTPNKKAQAQSAIESPLNFKDDDTHEGRKNTS

4. DSEENVQVQA(P)TPAVKKRTVTPNKKAQAQSAIE(P)SPLNFKDDDTHEGRKNTS

R-mutant (peptides 3 and 4)

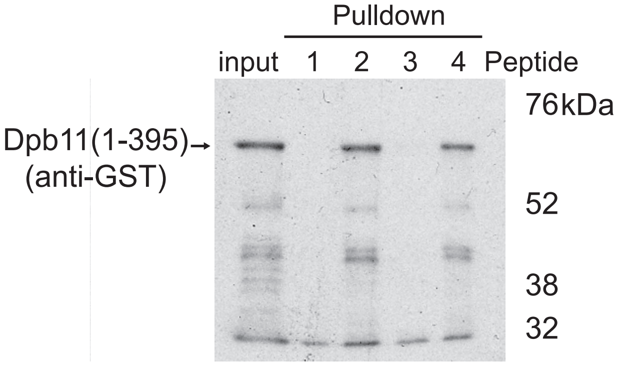


C

| Genotype | Viable |
|---|---|
| *sld3-R orc2-1* | yes |
| *sld3-R cdc6-1* | yes |
| *sld3-R 3HA-sld5 (hypomorph)* | yes |
| *sld3-R sml1Δ rad53Δ* | yes |

D

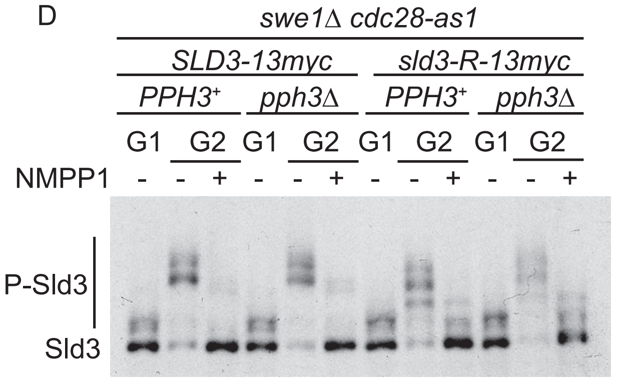


E

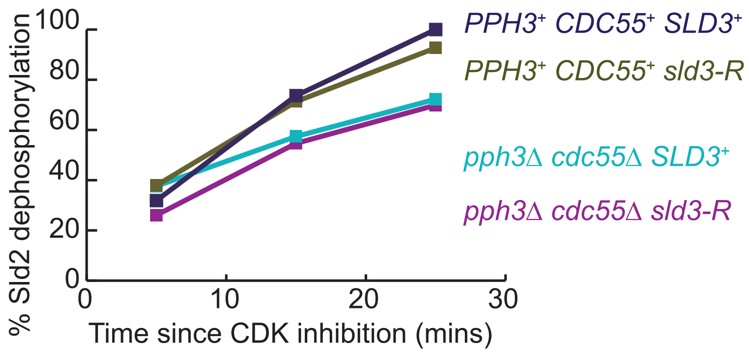


F

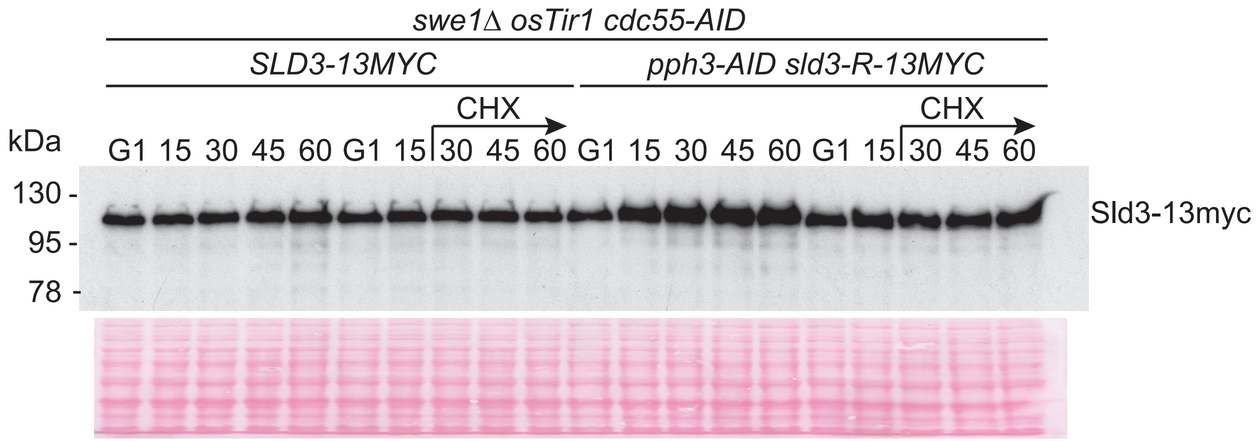

**Supplementary Figure 5.**

A. Alignment of Sld3 from yeast species, indicating the conservation of the essential CDK sites (red arrows), T600 and S622 in *S. cerevisiae*, as well as the Rts1 binding consensus sequence LxxIxE. The Rts1 binding motif consensus also indicates a preference for basic residues in the preceding amino acids [3], indicated here with blue arrows.

B. Top: Peptides of Sld3 (590-639) used to analyse Dpb11 binding. Bottom: anti-GST western blot of a peptide pulldown of a GST-Dpb11 fragment encompassing the first two BRCT repeats (1-395).

C. Viability of the *sld3-R* mutant (L616A, I619A) with other replication or checkpoint mutant strains, assessed by tetrad dissection of a heterozygous diploid.

D. Anti-myc phos-tag western blots of the indicated strains. Cells were either arrested in G1 phase or in G2 phase with nocodazole, with (+) or without (-) the addition of 1-NM-PP1 for 10 minutes.

E. Quantification of Sld2 dephosphorylation from main Figure 4F as in main Figure 3G.

F. Anti-myc western blot of Sld3-13myc on an SDS-PAGE gel (without phos-tag) from the indicated strains with and without addition of cycloheximide (CHX). A ponceau staining of this gel is shown below as a loading control.

Supplementary Figure 6.

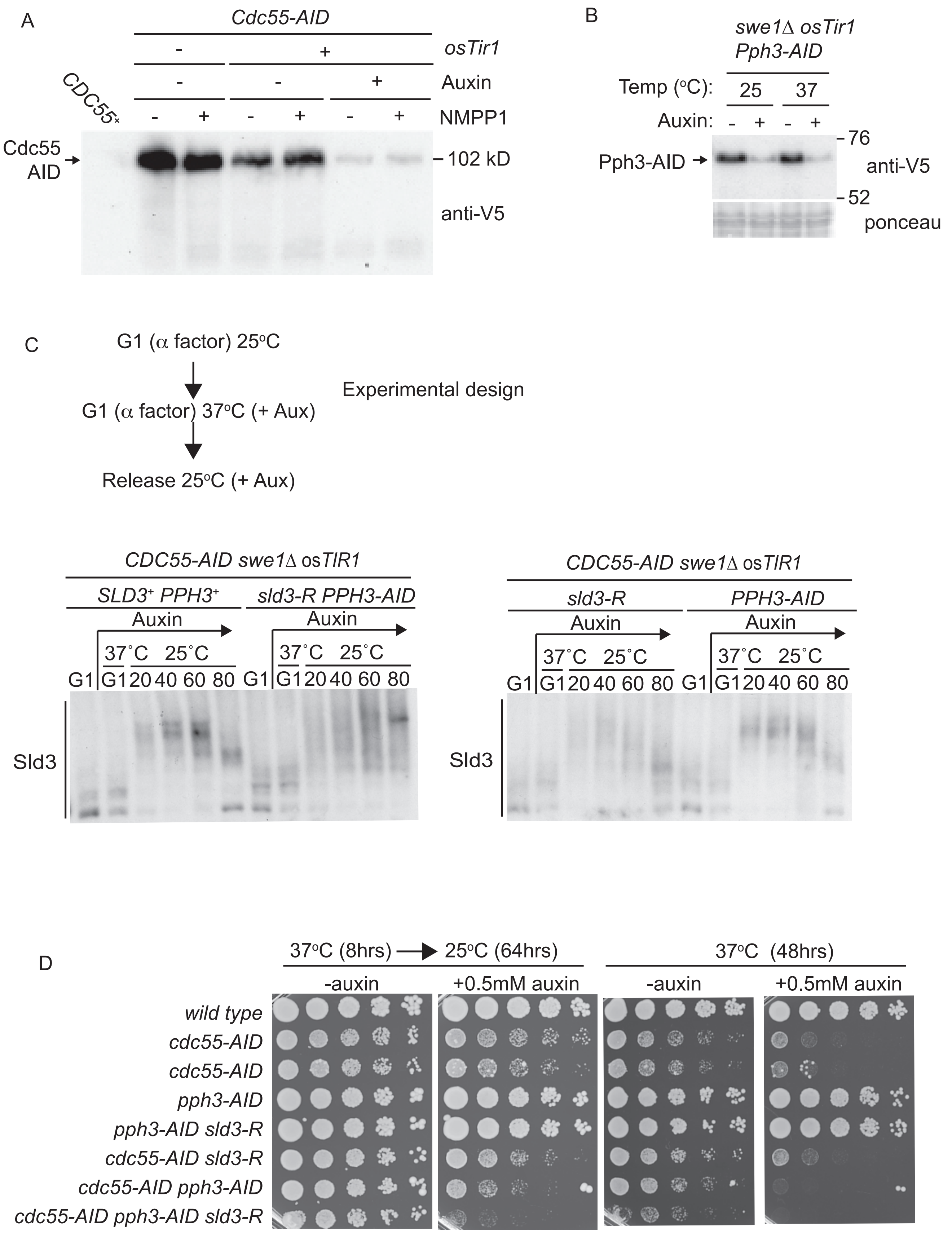

**Supplementary Figure 6.**

A. Anti-V5 western blot of the indicated strains with and without the addition of auxin for 30 minutes. Cdc55-AID contains a V5 tag.

B. As A.

C. Anti-myc western blot of Sld3-13myc, either wild type *SLD3*$^+$ or *sld3-R*, on a phos-tag SDS-PAGE gel. Cells were arrested and released according to the experimental design above.

D. Growth assay of the indicated strains. All strains are *swe1*Δ and express osTIR1. The 2 plates on the left were pre-grown at 37°C for 8 hours before switching to 25°C, similar to the method in C.

Supplementary Figure 7

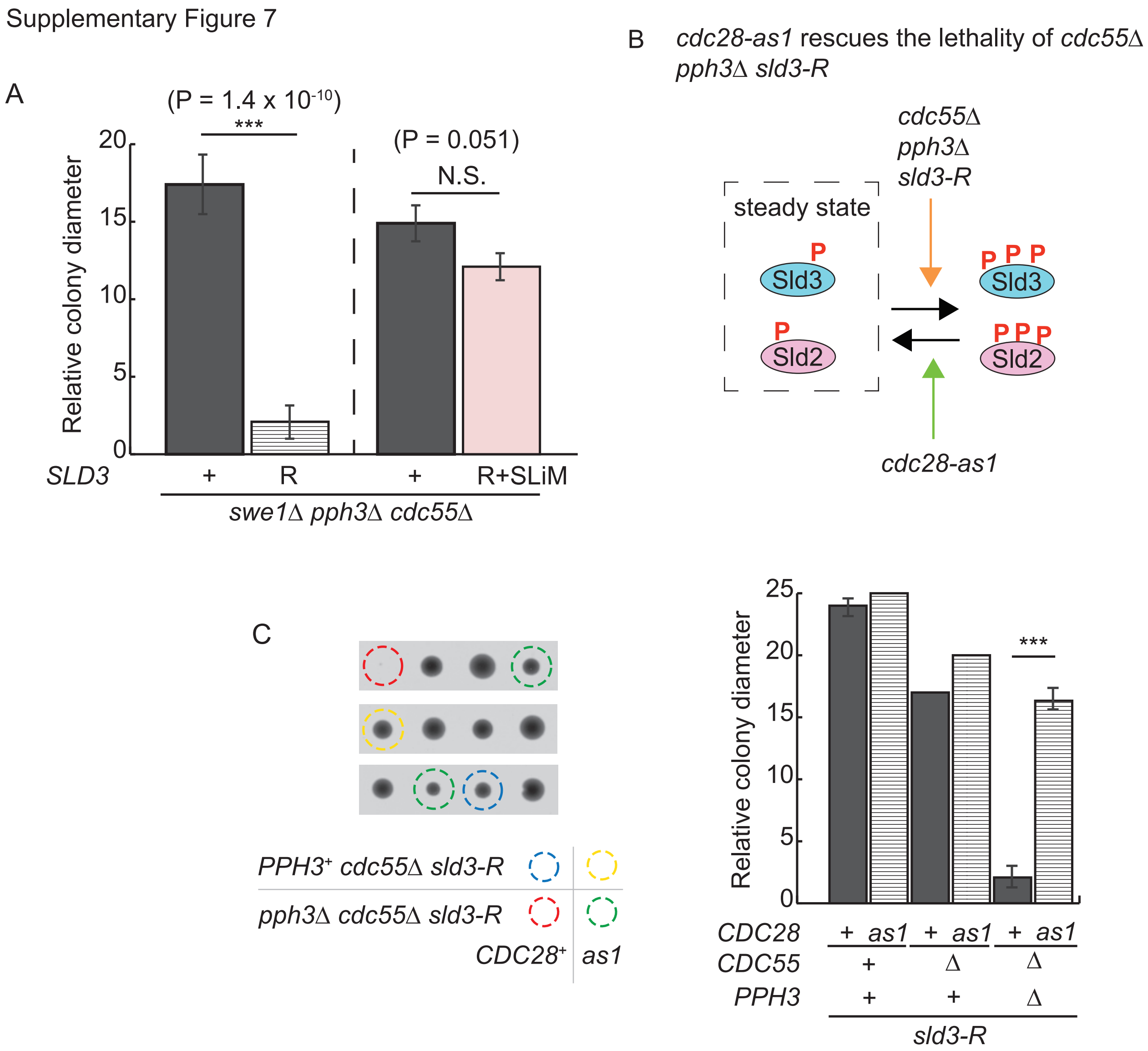


**Supplementary Figure 7.**

A. Relative spore colony diameter of the indicated strains with either wild type *SLD3* (+), *sld3-R* (R), or *sld3-R* with the Rts1 SLiM fused at the C-terminus (R+SLiM). All strains are *swe1*Δ. P values are from t-tests.

B. Diagram of how the different mutant backgrounds may affect the steady state level of Sld3/Sld2 phosphorylation, explaining why the hypomorphic *cdc28-as1* mutant partially rescues the phosphatase mutant phenotype in C.

C. Example tetrads (Left) and relative spore colony diameter (right) of the indicated strains. All strains are *swe1*Δ. *** refers to P value 0.005 from a t-test.

**Supplementary Figure references.**


1. Mattarocci, S. *et al.* Rif1 controls DNA replication timing in yeast through the PP1 phosphatase Glc7. *Cell reports* **7**, 62-69 (2014).

2. Kalev, P. & Sablina, A.A. Protein phosphatase 2A as a potential target for anticancer therapy. *Anti-cancer agents in medicinal chemistry* **11**, 38-46 (2011).

3. Wang, X. *et al.* A dynamic charge-charge interaction modulates PP2A:B56 substrate recruitment. *Elife* **9** (2020).
