## Supplemental Table 1 for "Dephosphorylation of the pre-initiation complex during S-phase is critical for origin firing"

Supplementary Table 1.

| Strain | Relevant genotype |
|---|---|
| W303a | *MATa ade2-1 ura3-1 his3-11,15 trp1-1 leu2-3,112 can1-100* |
| PZ1584 | *W303a cdc28-as1 (F88G) SLD3-10HIS-13MYC::KanMx* |
| PZ2474 | *W303a cdc28-as1 (F88G) SLD2-13MYC::KanMx* |
| PZ3253 | *W303a cdc28-as1 (F88G) YEN1-13MYC::KanMx* |
| PZ2280 | *W303a cdc28-as1 (F88G) SLI15-3HA::HIS3Mx* |
| PZ1854 | *W303a cdc28-as1 (F88G) Trp1::Sld3-10his-13myc::TRP1* |
| PZ3513 | *W303a cdc28-as1 (F88G) Trp1::Sld3-10A-10his-13myc::TRP1* |
| PZ3514 | *W303a cdc28-as1 (F88G) Trp1::Sld3-12A-10his-13myc::TRP1* |
| PZ52 | *W303a SLD3-10HIS-13MYC::KanMx* |
| PZ1694 | *W303a Sld3-10his13myc::KanMx clb5Δ::URA3* |
| PZ1699 | *W303a SLD2-13MYC::KanMx* |
| PZ1696 | *W303a clb5Δ::KanMx Sld2-13myc::KanMx* |
| PZ1198 | *W303a cdc28-as1 (F88G)* |
| PZ1322 | *W303a cdc28-as1 (F88G) cdc14-1::TRP1* |
| PZ3395 | *W303a cdc28-as1 (F88G) SLD3-10HIS-13MYC::KanMx swe1Δ::TRP1* |
| PZ3408 | *W303a cdc28-as1 (F88G) SLD2-13MYC::KanMx swe1Δ::TRP1* |
| PZ3457 | *W303a cdc28-as1 (F88G) SLD3-10HIS-13MYC::KanMx swe1Δ::TRP1 pph21Δ::HIS3 pph22Δ::HPHMx* |
| PZ3445 | *W303a cdc28-as1 (F88G) SLD2-13MYC::KanMx swe1Δ::TRP1 pph21Δ::HIS3 pph22Δ::HPHMx* |
| PZ2953 | *W303a SLD3-10HIS-13MYC::KanMx pph21Δ::HIS3 pph22Δ::HPHMx* |
| PZ3042 | *W303a SLD3-10HIS-13MYC::KanMx tpd3Δ::LEU2* |
| PZ3034 | *W303a SLD3-10HIS-13MYC::KanMx rts3Δ::LEU2* |
| PZ3102 | *W303a SLD3-10HIS-13MYC::KanMx cdc55Δ::LEU2* |
| PZ3075 | *W303a SLD3-10HIS-13MYC::KanMx rts1Δ::LEU2* |
| PZ3300 | *W303a SLD3-10HIS-13MYC::KanMx rts1Δ::LEU2 cdc55Δ::LEU2* |
| PZ3474 | *W303a cdc28-as1 (F88G) SLD3-10HIS-13MYC::KanMx swe1Δ::TRP1 tpd3Δ::LEU2* |
| PZ3650 | *W303a cdc28-as1 (F88G) SLD3-10HIS-13MYC::KanMx swe1Δ::TRP1 pph3Δ::HphNT* |
| PZ3561 | *W303a cdc28-as1 (F88G) SLD3-10HIS-13MYC::KanMx swe1Δ::TRP1 tpd3Δ::LEU2 pph3Δ::HphNT* |
| PZ3470 | *W303a cdc28-as1 (F88G) SLD2-13MYC::KanMx swe1Δ::TRP1 tpd3Δ::LEU2* |
| PZ3519 | *W303a cdc28-as1 (F88G) SLD2-13MYC::KanMx swe1Δ::TRP1 pph3Δ::HphNT* |
| PZ3559 | *W303a cdc28-as1 (F88G) SLD2-13MYC::KanMx swe1Δ::TRP1 tpd3Δ::LEU2 pph3Δ::HphNT* |
| PZ3623 | *W303a cdc28-as1 (F88G) trp1::sld3-10A-10his-13myc::TRP1 swe1Δ::TRP1 tpd3Δ::LEU2* |
| PZ3625 | *W303a cdc28-as1 (F88G) trp1::sld3-10A-10his-13myc::TRP1 swe1Δ::TRP1 pph3Δ::HphNT* |
| PZ3712 | *W303a cdc28-as1 (F88G) trp1::sld3-10A-10his-13myc::TRP1 swe1Δ::TRP1 tpd3Δ::LEU2 pph3Δ::HphNT* |
| PZ3713 | *W303a cdc28-as1 (F88G) trp1::sld3-12A-10his-13myc::TRP1 swe1Δ::TRP1 tpd3Δ::LEU2 pph3Δ::HphNT* |
| PZ2365 | *L40/YHGX13 diploid, gal4::loxP-kanMX-loxP/gal4Δ ade2 trp1-901/ade2-101::loxP-kanMX-loxP,*<br>*leu2-3,112/ leu2-3,-112, his3delta200/ hisdelta 200, LYS2/LYS2::(lexAop)4-HIS3, ura3-52::URA3(lexAop)8-lacZ/ ura3-52 URA3::UASGAL1-LacZ* |
| PZ3601 | *W303a SLD3-13MYC::KanMx cdc28-as1 (F88G) swe1Δ::TRP1* |
| PZ3602 | *W303a sld3-R(2A)-13myc::KanMx cdc28-as1 (F88G) swe1Δ::TRP1* |

| | |
|---|---|
| PZ3597 | *W303a cdc55Δ::LEU2 SLD3-13MYC::KanMx cdc28-as1 (F88G) swe1Δ::TRP1* |
| PZ3598 | *W303a cdc55Δ::LEU2 sld3-R(2A)-13myc::KanMx cdc28-as1 (F88G) swe1Δ::TRP1* |
| PZ3632 | *W303a cdc55Δ::LEU2 SLD3-13MYC::KanMx cdc28-as1 (F88G) swe1Δ::TRP1 pph3Δ::HphNT* |
| PZ3656 | *W303a cdc55Δ::LEU2 sld3-R(2A)-13myc::KanMx cdc28-as1 (F88G) swe1Δ::TRP1 pph3Δ::HphNT* |
| PZ3714 | *W303a cdc28-as1 (F88G) swe1Δ::TRP1 trp1::sld3-10A-R(2A)-13myc::TRP1* |
| PZ3716 | *W303a cdc28-as1 (F88G) swe1Δ::TRP1 cdc55Δ::LEU2 pph3Δ::HphNT trp1::sld3-10A-R(2A)-13myc::TRP1* |
| PZ3717 | *W303a cdc28-as1 (F88G) cdc55Δ::LEU2 pph3Δ::HphNT trp1::sld3-12A-R(2A)-13myc::TRP1* |
| PZ3868 | *W303a cdc28-as1 (F88G) swe1Δ::TRP1 SLD2-13MYC::KanMX SLD3-R(2A)::HIS3MX* |
| PZ3890 | *W303a cdc28-as1 (F88G) swe1Δ::TRP1 SLD2-13MYC::KanMX SLD3-R(2A)::HIS3MX cdc55Δ::LEU2 pph3Δ::HphMX* |
| PZ3888 | *W303a cdc28-as1 (F88G) swe1Δ::TRP1 SLD2-13MYC::KanMX pph3Δ::HphMX cdc55Δ::LEU2* |
| PZ3911 | *W303a swe1Δ::TRP1 cse4Δ::NatMX leu2::CSE4-HA6::LEU2 CDC55-AID::KanMx his3::PGPD1-osTir1::HIS3 SLD3-13myc::KanMx Mcm4-3HA::KanMX* |
| PZ3905 | *W303a swe1Δ::TRP1 cse4Δ::NatMX leu2::CSE4-HA6::LEU2 CDC55-AID::KanMx his3::PGPD1-osTir1::HIS3 sld3-R(2A)-13myc::KanMx Mcm4-3HA::KanMX* |
| PZ3942 | *W303a swe1Δ::TRP1 cse4Δ::NatMX leu2::CSE4-HA6::LEU2 CDC55-AID::KanMx PPH3-AID::HphMx his3::PGPD1-osTir1::HIS3 SLD3-13myc::KanMx Mcm4-3HA::KanMX* |
| PZ3966 | *W303a swe1Δ::TRP1 cse4Δ::NatMX leu2::CSE4-HA6::LEU2 CDC55-AID::KanMx PPH3-AID::HphMx his3::PGPD1-osTir1::HIS3 sld3-R(2A)-13myc::KanMx Mcm4-3HA::KanMX* |
| PZ4450 | *W303a swe1Δ::TRP1 sld3-13myc::KanMx his3::PGPD1-osTir1::HIS3 CDC55-AID::KanMx* |
| PZ4469 | *W303a swe1Δ::TRP1 sld3-R(2A)-13myc::KanMx his3::PGPD1-osTir1::HIS3 PPH3-AID::HphMx CDC55-AID::KanMx* |
| PZ3998 | *W303a swe1Δ::TRP1 cse4Δ::NatMX leu2::CSE4-HA6::LEU2 CDC55-AID::KanMx his3::PGPD1-osTir1::HIS3 sld3-13myc::KanMx Sld2-6HA::URA3* |
| PZ3987 | *W303a swe1Δ::TRP1 cse4Δ::NatMX leu2::CSE4-HA6::LEU2 CDC55-AID::KanMx PPH3-AID::HphMx his3::PGPD1-osTir1::HIS3 sld3-R(2A)-13myc::KanMx Sld2-6HA::URA3* |
| PZ3427 | *W303alpha cdc55Δ::LEU2 swe1Δ::TRP1* |
| PZ4003 | *W303a swe1Δ::TRP1 SLD3-R(2A)::HIS3MX pph3Δ::HphMX* |
| PZ4059 | *W303a swe1Δ::TRP1 Sld3-R(2A)-SLIM-13myc::KanMX* |
| PZ3641 | *W303alpha cdc55Δ::LEU2 swe1Δ::TRP1 pph3Δ::HphNT* |
| PZ4133 | *W303a YOR131C::PGPD1-GAL4-ER-CYC1 terminator::NatMX::YOR131C SLD3-R(2A)-13MYC::KanMx swe1Δ::TRP1 leu2::PGPD1-OsTIR1::LEU2 CDC55-AID::KanMx pph3Δ::HphNT* |
| PZ4136 | *W303a YOR131C::PGPD1-GAL4-ER-CYC1 terminator::NatMX::YOR131C SLD3-R(2A)-13MYC::KanMx swe1Δ::TRP1 leu2::PGPD1-OsTIR1::LEU2 CDC55-AID::KanMx pph3Δ::HphNT his3::PGAL-sld3-R(2A)::HIS3 ura3::PGAL-Dpb11::URA3* |
| PZ4101 | *W303a YOR131C::PGPD1-GAL4-ER-CYC1 terminator::NatMX::YOR131C SLD3-R(2A)-13MYC::KanMx swe1Δ::TRP1 leu2::PGPD1-OsTIR1::LEU2 CDC55-AID::KanMx ura3::PGAL-Dpb11::URA3* |

| | |
|---|---|
| PZ4091 | *W303alpha YOR131C::PGPD1-GAL4-ER-CYC1 terminator::NatMX::YOR131C his3::PGAL-sld3-R(2A)::HIS3 SLD3-R(2A)-13MYC::KanMx swe1Δ::TRP1 leu2::PGPD1-OsTIR1::LEU2 pph3Δ::HphNT* |
| PZ3277 | *W303a cdc28-as1 (F88G) Trp1::Sld2-13Myc::TRP1MX* |
| PZ3278 | *W303a cdc28-as1 (F88G) Trp1::Sld2-T84A-13Myc::TRP1MX* |
| PZ2014 | *W303a cdc28-as1 (F88G) swe1Δ::TRP1 ura3::PSLD2-SLD2-13myc::URA3* |
| PZ2015 | *W303a cdc28-as1 (F88G) swe1Δ::TRP1 ura3::PSLD2-sld2-3D3A-13myc::URA3* |
| PZ2016 | *W303a cdc28-as1 (F88G) swe1Δ::TRP1 ura3::PSLD2-sld2-3D4A-13myc::URA3* |
| PZ2017 | *W303a cdc28-as1 (F88G) swe1Δ::TRP1 ura3::PSLD2-sld2-3D7A-13myc::URA3* |
| PZ2018 | *W303a cdc28-as1 (F88G) swe1Δ::TRP1 ura3::PSLD2-sld2-3D8A-13myc::URA3* |
| PZ3434 | *W303a cdc28-as1 (F88G) trp1::PSLD2-sld2-10D-13myc::TRP1* |
| PZ3436 | *W303a cdc28-as1 (F88G) trp1::PSLD2-sld2-11D-13myc::TRP1* |
| PZ1850 | *W303a cdc6-1 cdc28-as1 (F88G) Sld3-10his13myc::KanmX* |
| PZ869 | *W303a Sld3-3xFlag::hygro* |
| PZ1388 | *W303a Sld3-3xFlag::hygro cdc28-as1 (F88G)* |
| PZ1355 | *W303a CDC14-1::TRP1 cdc28-as1 (F88G) Sld3-3xFlag::hygro* |
| PZ1763 | *W303a cdc28-as1 (F88G) Sld3-10his13myc::KanMx rif1Δ::URA3* |
| PZ2097 | *W303a pdr5Δ::hphMX6 cdc28-as1 (F88G) Sld3-10his-13myc::KanMx* |
| PZ3433 | *W303a cdc28-as1 (F88G) SLD3-10HIS-13MYC::KanMx rts1Δ::LEU2 cdc55Δ::LEU2 swe1Δ::TRP1* |
| PZ3431 | *W303a cdc28-as1 (F88G) SLD2-13MYC::KanMx rts1Δ::LEU2 cdc55Δ::LEU2 swe1Δ::TRP1* |
| PZ623 | *W303alpha orc2-1* |
| PZ3606 | *W303a sld3-R(2A)-13myc::KanMx swe1Δ::TRP1* |
| PZ209 | *W303alpha cdc6-1* |
| PZ2481 | *W303alpha 3HA-sld5::KanMx* |
| PZ3422 | *W303a cdc55Δ::LEU2 SLD2-13MYC::KanMx swe1Δ::TRP1* |
| PZ3897 | *W303a swe1Δ::TRP1 SLD2-13MYC::KanMX SLD3-R(2A)::HIS3MX pph3Δ::HphMX* |
| PZ3664 | *W303a sld3-R(2A)-13myc::KanMx cdc28-as1 (F88G) swe1Δ::TRP1 pph3Δ::HphNT* |
| PZ3650 | *W303a SLD3-13MYC::KanMx cdc28-as1 (F88G) swe1Δ::TRP1 pph3Δ::HphNT* |
| PZ254 | *W303a pph3Δ::HphNT* |
| PZ3614 | *W303alpha cdc55Δ::LEU2 sld3-R(2A)-13myc::KanMx cdc28-as1 (F88G) swe1Δ::TRP1* |
| PZ3585 | *W303a swe1Δ::TRP1 cse4Δ::NatMX leu2::CSE4-HA6::LEU2 CDC55-AID::KanMx his3::PGPD1-osTir1::HIS3* |
| PZ3916 | *W303a swe1Δ::TRP1 cse4Δ::NatMX leu2::CSE4-HA6::LEU2 his3::PGPD1-osTir1::HIS3 SLD3-13myc::KanMx pph3-AID::HphMX* |
| PZ3918 | *W303a swe1Δ::TRP1 cse4Δ::NatMX leu2::CSE4-HA6::LEU2 his3::PGPD1-osTir1::HIS3 pph3-AID::HphMX sld3-R(2A)-13myc::KanMx* |
| PZ3924 | *W303a swe1Δ::TRP1 cse4Δ::NatMX leu2::CSE4-HA6::LEU2 his3::PGPD1-osTir1::HIS3 sld3-R(2A)-13myc::KanMx cdc55-AID::KanMX* |
| PZ3919 | *W303a swe1Δ::TRP1 cse4Δ::NatMX leu2::CSE4-HA6::LEU2 his3::PGPD1-osTir1::HIS3 SLD3-13myc::KanMx pph3-AID::HphMX cdc55-AID::KanMX* |
| PZ3922 | *W303a swe1Δ::TRP1 cse4Δ::NatMX leu2::CSE4-HA6::LEU2 his3::PGPD1-osTir1::HIS3 pph3-AID::HphMX sld3-R(2A)-13myc::KanMx cdc55-AID::KanMX* |
